# Geomorphic evolution of a Caribbean biological hotspot

**DOI:** 10.64898/2026.02.24.707483

**Authors:** Aaron O’Dea, Max Titcomb, H. Luke Anderson, Brígida de Gracia, Suzette G.A. Flantua, Michael G. Hynes, Thomas Parsons, Carmen Schlöder, Michael J. Braun

## Abstract

Continental shelf archipelagos offer rare opportunities to test island biogeography in a time-resolved, non-equilibrium setting because postglacial sea-level rise drives datable changes in both island area and isolation. Yet most near-shore systems still lack reconstructions that integrate spatial changes in island configuration with duration-weighted histories of fragmentation. Such information is essential for interpreting modern genomic and biodiversity patterns and for evaluating responses to fragmentation such as extinction debt, immigration lag and adaptive responses over time. Here we reconstruct the land-sea geomorphic history of the Bocas del Toro Archipelago, Panama, combining high-resolution bathymetry and topography with a sea-level model corrected for sedimentation and tectonic movement. From this, we quantify multi-scale isolation, historical fragmentation dynamics, and composite metrics that integrate space and time. We show that present-day islands became isolated sequentially from 9.5 to 2.9 ka, with post-isolation area contraction rates varying 20-fold among islands. Over the same interval, shallow marine habitat (0-10 m depth) expanded ninefold as shelf flooding created space for reefs, seagrass, and mangroves. Extending the analyses through the Pleistocene reveals the modern archipelago is atypical and for much of the last million years, the region existed as continuous coastal lowland rather than discrete islands. Projections under moderate emissions scenarios predict ∼5% terrestrial habitat loss but ∼50% expansion of shallow marine habitat by 2150, though whether degraded Caribbean reefs can exploit this expansion remains uncertain. Exploratory correlations between species richness of terrestrial vertebrate groups and geomorphometric predictors reveal that island area, maximum elevation, and cumulative habitat availability since isolation are the strongest correlates of diversity, whilst buffer-zone isolation indices are confounded by covariation between island size and surrounding landmass in this continental shelf context. Overall, Bocas del Toro provides a time-calibrated natural experiment for testing how dynamic connectivity and isolation shaped island and coastal biodiversity.

## Introduction

Archipelagos located close to large land masses provide tractable systems for exploring how geomorphic dynamics drive ecological and evolutionary processes over millennial timescales. In these systems, insular fragments become isolated during late Pleistocene and Holocene sea level rise creating well-defined histories of fragmentation. Unlike oceanic island systems that typically emerge through volcanism, these “land-bridge” or “stepping stone” archipelagos (Flantua et al. 2020; Itescu et al. 2020) begin as continuous coastal lowlands during sea level lowstands that may harbor the full complement of mainland biodiversity. Rising seas then drive sequential isolation of islands, creating clear spatial boundaries and knowable fragmentation histories (Rijsdijk et al. 2014; Simaiakis et al. 2017; Ávila et al. 2025). Such islands may initially capture a large fraction of mainland biodiversity that decays to lower levels over time, driven by the ecological and evolutionary processes that govern species-area relationships (Matthews et al. 2021). The same sea level rise that fragments terrestrial habitats simultaneously transforms adjacent shallow marine ecosystems, which in tropical areas often consist of highly diverse reef, seagrass and mangrove habitats. As rising seas inundate continental shelves, accommodation space for marine systems tends to expand, in sharp contrast to the accompanying loss of land. The timing and magnitude of these changes depend on the dynamic interplay between local bathymetric configuration, tectonics, sediment infilling and sea level change.

Despite this geomorphic coupling between land and sea, island biogeography has traditionally focused on terrestrial systems, suffering from what Pinheiro et al. (2017) identified as a “historical imbalance” wherein marine organisms have received comparatively little attention despite occupying environments equally shaped by sea level fluctuations and of similar ecological significance. In terrestrial island biogeography, advances over the last 15 years have demonstrated that simple distance-to-mainland metrics fail to capture spatial complexity (Itescu et al. 2020). Isolation indices accounting for surrounding landmass proportions across multiple spatial scales explain substantially more variation in species richness than distance alone (Weigelt & Kreft 2013), whilst buffer-based approaches incorporating stepping-stone connectivity better predict β-diversity patterns (Itescu et al. 2020). These findings reveal that colonisation and gene flow operate across multiple spatial scales simultaneously, from local stepping-stones to regional mainland source pools.

Temporal dynamics are equally important. Intermediate sea level configurations persist far longer than extremes like the Last Glacial Maximum (LGM) and may play a larger role in shaping biodiversity patterns, underscoring the importance of duration-weighted histories rather than snapshot analyses (Ali & Aitchison 2014; Norder et al. 2019; Flantua et al. 2020). Yet whether historical configurations predict contemporary diversity varies: past mainland connections structured richness across Aegean shelf islands (Hammoud et al. 2021), whereas Sundaic bird diversity equilibrated rapidly after post-LGM isolation (Sin et al. 2022). Phylogenetic analyses confirm that colonisation rates decline with isolation, extinction rates decline with area, and speciation rates increase with both (Valente et al. 2020). Reconstructing geomorphic histories that quantify fragmentation timing, area contraction, and duration-weighted configurations facilitates tests of such theory (Pinheiro et al. 2017; Ávila et al. 2019).

The Bocas del Toro Archipelago on Panama’s Caribbean coast offers an excellent system for developing such an integrated framework. The region sits on the Panamá Volcanic Arc, which formed from Cocos Ridge subduction beginning in the Miocene, driving regional uplift that contributed to Isthmus of Panama emergence around 3 million years ago (O’Dea et al. 2016). During the Last Glacial Maximum, present-day islands formed a contiguous coastal plain connected to the mainland. Post-glacial sea level rise progressively fragmented this landscape, sequentially isolating islands at different times based on coastal topography and shelf relief (Titcomb and O’Dea 2020). The region supports high biodiversity across both realms: lowland tropical forest and peat swamps on land; extensive coral reefs, seagrass beds, and mangrove forests in the sea (Guzman et al. 2005; Sjögersten et al. 2011; Seemann et al. 2018; Collin et al. 2024). Rapid evolutionary responses to isolation have already been documented, including dramatic colour polymorphism in poison frogs, hybridisation-driven trait introgression in manakins, insular gigantism in birds, and insular dwarfism in sloths (Wetmore 1959, 1963; Parsons et al. 1993; Anderson & Handley Jr 2002; Wang & Shaffer 2008; Gehara et al. 2014). Reconstructing these landscape trajectories through the late Pleistocene and Holocene also provides useful archaeological context, as the availability of terrestrial and marine habitats to pre-European human coastal settlement would have depended greatly on the changing configurations of terrestrial and marine habitats (Cybulski et al. 2025).

Despite this rich biological and human context, comprehensive geomorphometric characterisation integrating spatial configuration and temporal dynamics has been lacking. Global bathymetric models derived from satellite altimetry, whilst valuable for broad-scale patterns (Norder et al. 2019; De Groeve et al. 2022), are resolution-limited in coastal settings and valid bathymetries fall below 10% near coasts (Cazenave et al. 2022) and island slope soundings commonly have uncertainties of 150–400 m (Klein et al. 2023). These limitations have direct biogeographical implications; accurate local bathymetries can adjust land bridge connection timings by thousands of years (Garg et al. 2022). Small-scale surveys and in-situ depth measurements are therefore essential for archipelago-scale reconstructions.

Here we reconstruct the integrated geomorphological history of the Bocas del Toro Archipelago, combining fine-scale bathymetric and topographic data with sea level curves corrected for sediment accumulation and tectonic movement. We calculate modern isolation metrics following established frameworks (Weigelt & Kreft 2013; Itescu et al. 2020), historical parameters quantifying fragmentation dynamics, and composite metrics integrating spatial and temporal dimensions. Whether these indices perform equivalently in continental shelf archipelagos remains largely untested. We simultaneously quantify shallow marine habitat dynamics through the last 12 kyr, place these within the Pleistocene context (last 1 Myr), and project changes to 2150 under continued sea level rise. To demonstrate the framework’s utility, we present exploratory analyses correlating species richness of five terrestrial vertebrate groups from Smithsonian museum collections with geomorphometric predictors.

### Methodology and approach

Our geomorphometric framework involves numerous analytical steps. We therefore present an abbreviated methodology here, with full procedural detail in Supplementary Methods. Table 1 defines all 15 major geomorphometric metrics and their calculations. Table S1 reports values for all 14 focal islands. All code and data are available in the Zenodo repository (https://doi.org/10.5281/zenodo.18732423).

**Table 1.**
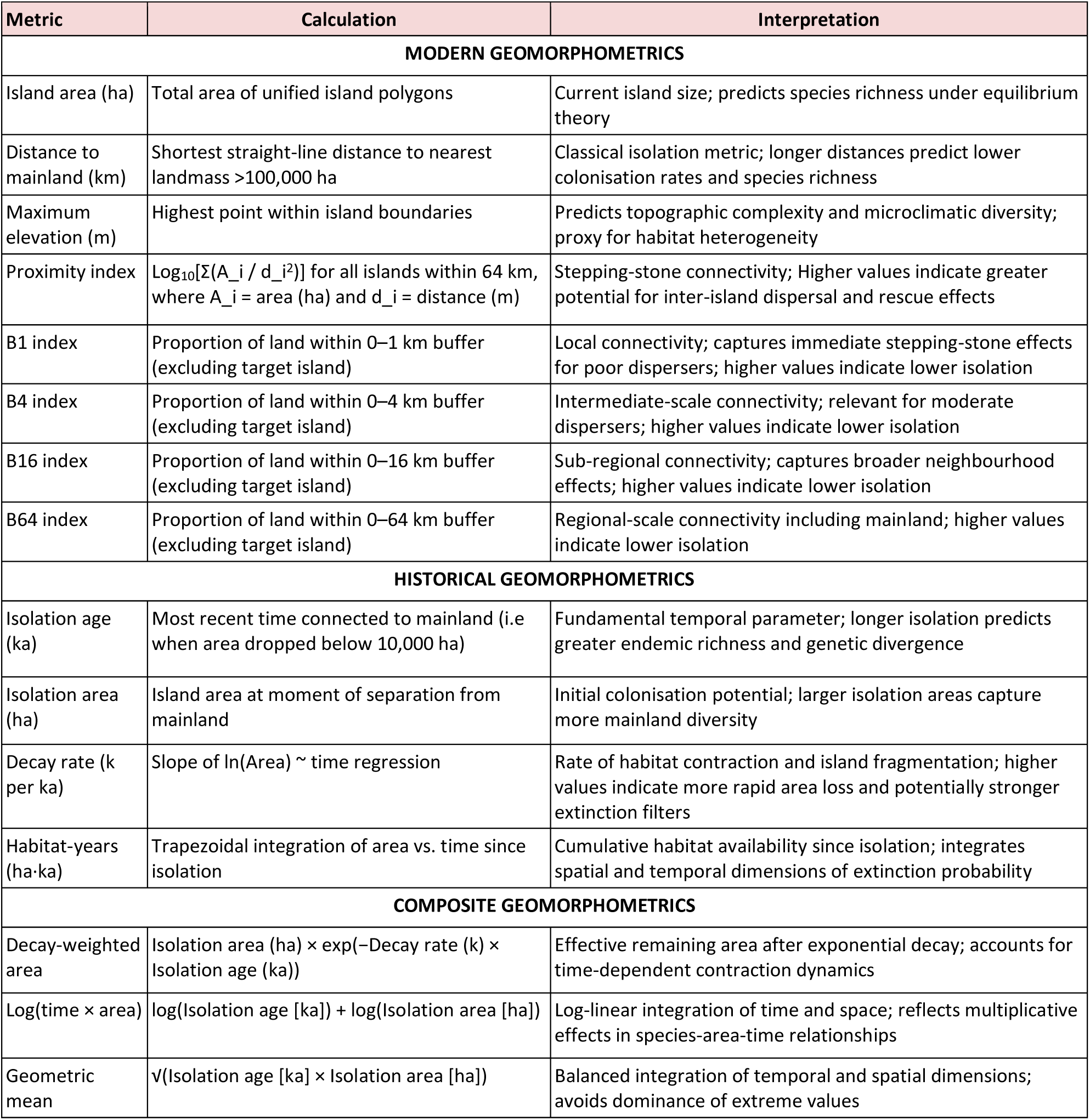
Geomorphometric metrics calculated for islands in the Bocas del Toro Archipelago, organised into modern spatial metrics (current configuration), historical metrics (temporal dynamics), and composite metrics (integrating spatial and temporal dimensions). Modern metrics follow established frameworks (Kalmar & Currie 2006; Weigelt & Kreft 2013; Itescu et al. 2020). Buffer indices increase with surrounding land area, thus higher B-index values indicate *lower* isolation. Historical metrics quantify post-glacial fragmentation dynamics from 12 ka to present. Composite metrics test alternative hypotheses about how isolation history structures contemporary biodiversity: exponential decay emphasises demographic collapse, log(time × area) emphasises habitat-time integration, and geometric mean provides neutral balance. See Methodology and Approach for detailed calculation procedures.

### Study area and digital elevation model

The Bocas del Toro Archipelago (Fig. 1; approximately 9°N, 82°W) lies along the Caribbean coast of northwestern Panama, comprising 14 major islands and hundreds of mangrove islets. The archipelago sits on the Panamá Arc, formed from the subduction of the Cocos Ridge beneath the Caribbean Plate beginning in the Miocene(Coates et al. 2003, 2005). The region experiences opposing tectonic forces: regional uplift from Cocos Ridge subduction versus episodic localised subsidence from earthquake-related thrust faulting (Plafker & Ward 1992; Suárez et al. 1995). The modern archipelago supports high terrestrial and marine biodiversity across lowland tropical wet forests, peat bogs, coral reefs, seagrass beds, and mangrove forests, making it a focal site for ecological and evolutionary research over the past three decades (Olson 1993; Guzman et al. 2005; Seemann et al. 2018; Collin et al. 2024). Archaeological evidence suggests humans arrived on the Isthmus between 25 and 16 ka (Ranere & Cooke 2021; Cybulski et al. 2025).

**Figure 1.**
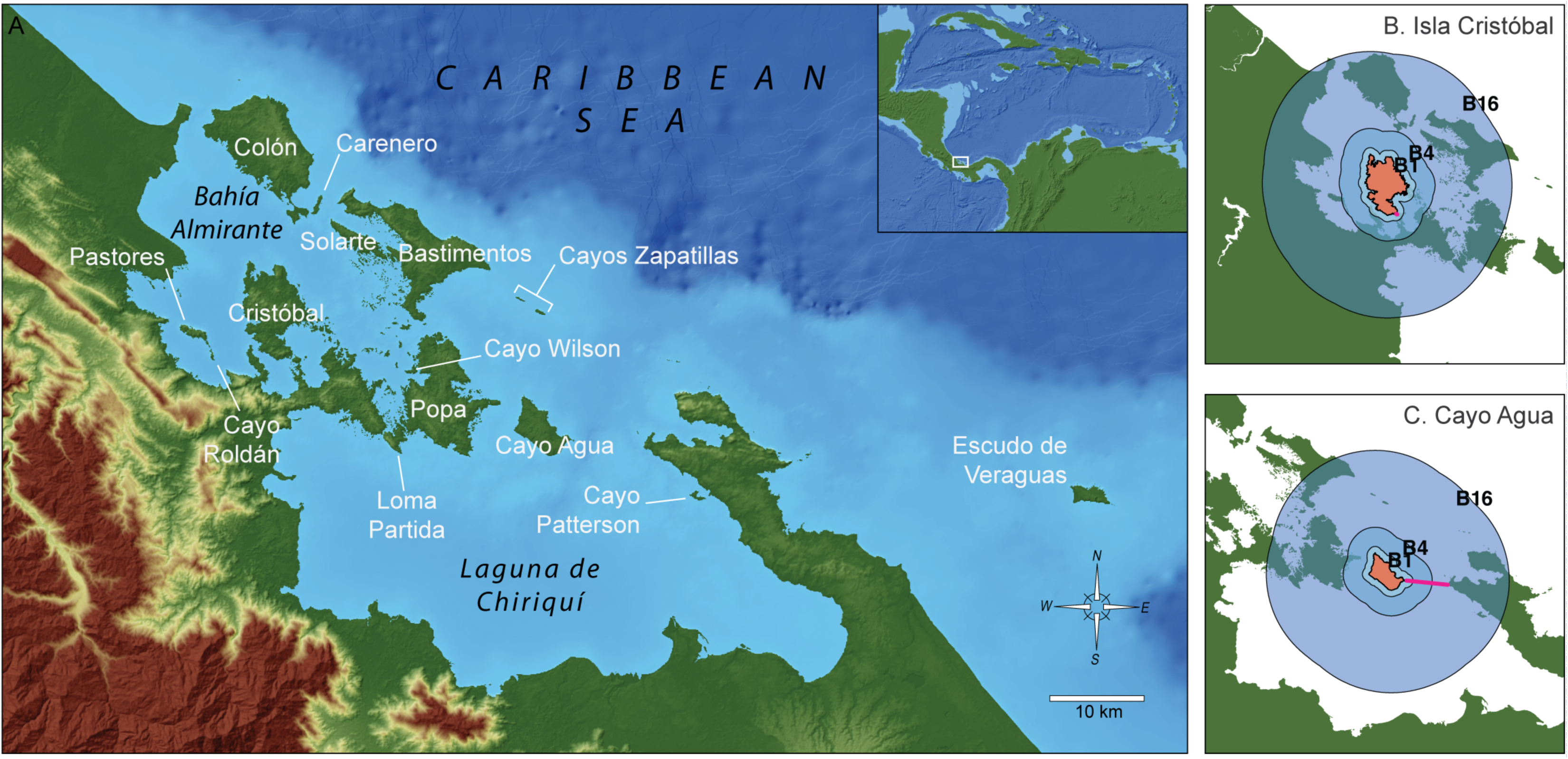
The Bocas del Toro Archipelago, Caribbean Panama. (A) The 14 principal islands and main water bodies. Inset shows the location of the archipelago in Tropical America. Light blue shading indicates bathymetry from 0 to 125 m. See Fig. S1 for a reconstruction of the region at the Last Glacial Maximum. (B, C) Buffer zone isolation analysis for two contrasting islands. Buffer zones B1, B4 and B16 (blue shading; B64 omitted for clarity) surround each target island (orange). Magenta lines indicate the shortest distance to the mainland. Isolation indices are calculated as the proportion of each buffer zone occupied by land (excluding the target island itself), with higher values indicating lower isolation. (B) Isla Cristóbal shows high buffer indices due to substantial mainland and adjacent island coverage within buffer zones. (C) Cayo Agua shows lower buffer indices with less surrounding land.

We created an integrated terrestrial-marine digital elevation model (DEM) by combining bathymetric data digitised from 14 high-resolution nautical charts (US Army Map Service Series E762, 1:25,000 to 1:50,000 scale) with NASA SRTM 30 m topography (Jarvis et al. 2008). All soundings were standardised to metres below present mean sea level and interpolated using Inverse Distance Weighting at 30 m resolution. Six channels with known field-measured depths were manually corrected to prevent spurious land bridges (see Supplementary Methods).

### Geomorphometric framework

We calculated 15 metrics characterising island configuration and isolation history, organised into modern spatial metrics, historical temporal metrics, and composite metrics integrating both dimensions (Table 1).

Modern metrics quantify current archipelago configuration. Island area, maximum elevation, and distance to mainland follow standard approaches. We calculated a proximity index following Kalmar & Currie (2006) as the log-transformed inverse-distance-weighted sum of surrounding island areas, and multi-scale buffer zone isolation indices (B1, B4, B16, B64) following Weigelt and Kreft (2013) and Itescu et al. (2020), representing the proportion of land within cumulative buffers at 1, 4, 16, and 64 km from each island (Fig. 1B,C). Higher buffer values indicate lower isolation.

Historical metrics quantify post-glacial fragmentation dynamics. We reconstructed island configurations at 100-year intervals through the Holocene (12 ka to present) by applying a Caribbean-specific relative sea level curve to our DEM, corrected for sediment accumulation and net tectonic movement (see Supplementary Methods). At each time step, eight-directional connectivity analysis identified contiguous land patches, allowing us to track island area through time, isolation from the mainland (defined as the largest contiguous landmass >100,000 ha), and post-isolation area decay. From these reconstructions we derived four historical metrics (Table 1): isolation age, isolation area at the moment of separation, decay rate (exponential area loss following Sin et al. 2022), and cumulative habitat-years (trapezoidal integration of area through time).

Composite metrics integrate temporal and spatial dimensions to test alternative hypotheses about how isolation history structures biodiversity. We calculated three composites (Table 1): decay-weighted area (emphasising recent effective area; Weigelt & Kreft 2013; Whittaker et al. 2017), log(time × area) (emphasising multiplicative habitat-time availability; Weigelt et al. 2016), and geometric mean (providing balanced integration without strong assumptions about relative importance).

### Holocene reconstructions

We developed a Caribbean-specific relative sea level curve by fitting a generalised additive model (GAM) to Caribbean sea level data from Lambeck et al. (2014) following Khan et al. (2017), constrained to pass through 0 m at present (Fig. S2). Full model in Supplementary Methods.

We estimated the area of potential coral reef, seagrass, and mangrove habitat through time by identifying seafloor at 0-10 m below contemporaneous sea level at each time step.

We calculated pairwise isolation ages between all islands and between each island and the mainland using connectivity analysis on historical land masks. This captures sequential fragmentation: islands that separated from the mainland as part of larger landmasses could continue exchanging populations until their later separation from each other. See Supplementary Methods.

### Temporal context

To assess whether modern configurations are typical or unusual, we extended analyses to the last million years using the global sea level curve of Clark et al. (2025). We sampled 1001 time points and calculated island number and total area at each, generating frequency distributions revealing which configurations persisted longest. This analysis provides lower resolution than the Holocene reconstruction.

We projected future habitat changes under a 2150 sea level rise scenario combining IPCC AR6 SSP3-7.0 estimates (+1.34 m; IPCC 2021) with predicted regional tectonic subsidence, yielding a total relative sea level rise of +1.64 m by 2150.

### Biogeographic analyses

Museum specimen records for Bocas del Toro were collated from collections housed at the US National Museum of Natural History (USNM) comprising mammals, herpetofauna, and birds (n = 11,378 after expert review). Records were standardised for island locality and taxonomy and consolidated into a dataset representing 624 unique species following procedures in Supplementary Methods.

To provide an initial demonstration of how the geomorphometric framework can inform biological research, we examined correlations between species richness and island geomorphometrics using the museum specimen dataset. We restricted analyses to five terrestrial vertebrate groups spanning a range of dispersal capabilities: Anura, resident non-migratory terrestrial birds, Chiroptera, Rodentia, and Squamata. For birds, migratory species and water birds were excluded using an expert-curated lookup table.

For each group, we calculated species richness per island, including only island-group combinations represented by five or more specimens to reduce the impact of incidental records on richness estimates. For each group, we computed Pearson correlation coefficients between species richness and each of 15 geomorphometric predictors (Table 1). We used simple correlations rather than multivariate regression because sample sizes were small (7 to 12 islands). Collecting effort varied substantially across islands and groups (Fig. S8), and these analyses should be considered exploratory.

## Results and Discussion

### Modern geomorphology and isolation metrics

The 14 principal islands range in area from 22 ha (Cayo Roldan) to 6,012 ha (Isla Colón), spanning nearly three orders of magnitude (Table S1, Figure 1). The three largest islands (Colón, Popa, and Bastimentos) together account for over half of the total archipelago land area. Numerous additional smaller islands and mangrove islets exist throughout the archipelago; whilst not analysed as focal islands, these were included in connectivity calculations as potential stepping stones.

Maximum elevations range from 14 m (Cayo Patterson) to 116 m (Loma Partida), reflecting underlying geology of the Miocene-Pliocene Panama Arc volcanic complex (Coates et al. 2003). Distance to mainland varies nearly 200-fold, from 80 m (Loma Partida) to 17.3 km (Escudo de Veraguas), providing substantial variation for testing classic island biogeographic predictions (MacArthur & Wilson 2001).

Simple distance metrics, however, capture only one dimension of spatial complexity. Our buffer-zone isolation indices reveal that islands at similar mainland distances can have very different surrounding landmass configurations (Fig. 1B,C, Table S1). For example, Escudo de Veraguas shows consistently high isolation across all spatial scales (B1–B16 indices all zero; B64 = 0.27), whilst Cristóbal shows low local connectivity (B1 = 0.03) but high regional connectivity (B64 = 0.41) due to its proximity to the mainland (Weigelt & Kreft 2013; Itescu et al. 2020).

The proximity index, which weights neighbouring island areas by inverse squared distance, varies from−5.4 (Escudo de Veraguas) to 1.8 (Cayo Wilson). This range spans seven orders of magnitude, reflecting the sensitivity of connectivity to nearby landmass configuration (Kalmar & Currie 2006) and potentially provides natural variation for testing how taxa with different dispersal capabilities respond to isolation. These modern metrics do however show varying degrees of intercorrelation (Fig. S3). As expected, buffer zone indices are, on the whole, all positively correlated with each other due to their nested spatial structure, whilst distance to mainland shows negative correlation with several connectivity metrics. However, most metrics capture sufficiently independent aspects of isolation to warrant their inclusion in biogeographic analyses.

### Holocene sea level reconstruction

Our Caribbean sea level model (Fig. S2) tracks the end of the rapid transgression from global glacial melt where sea level rose at an average of 9.4 mm/year between 12 and 8 ka. This rate slowed to 1.8 mm/year between 8 and 4 ka as sea levels began to stabilise. From 2 ka to the present sea level rise rate dropped further to 0.5 mm/year. These rates corroborate previous reconstructions (Toscano & Macintyre 2003; Khan et al. 2017) and provide the temporal framework for reconstructing island configurations throughout the Holocene.

### Terrestrial Holocene dynamics

At 12 ka, when sea level stood ∼60 m below present, the Bocas del Toro region formed a broad coastal plain connected to lowland forests extending across Caribbean Central America and into northwestern South America (Fig. 2). No landmass exceeding 20 ha existed as an island at this time (Fig. 2 & 3A). By 9 ka, outlines of the major islands had emerged, though many remained interconnected to each other or to the mainland. Progressive fragmentation continued through the mid-Holocene (8–6 ka) as rising seas invaded lower-elevation saddles between topographic highs.

**Figure 2.**
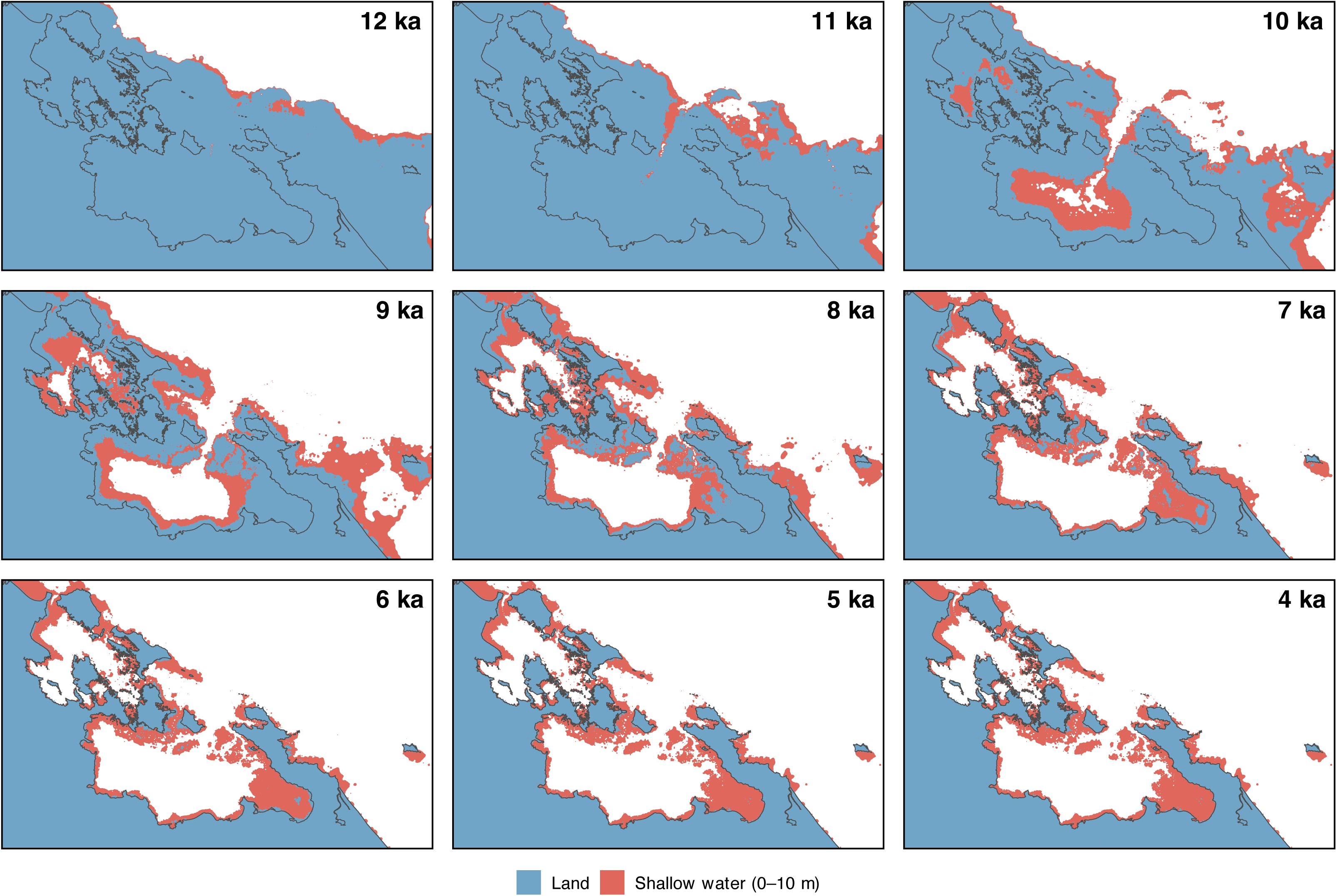
Formation of the Bocas del Toro Archipelago and its shallow water habitats from 12 to 4 ka. Blue areas show land above contemporary sea level, red areas the 0–10 m depth zone where coral reefs, seagrass and mangroves predominate. Reconstructions incorporate the regional sea level curve and account for channel sedimentation, tectonic uplift and subsidence at each time step. Sea level stabilised by ∼3 ka, so configurations after 4 ka are not reproduced. The thin dark grey line marks the modern coastline for orientation.

The number of islands larger than 20 ha peaked at 40 at 7 ka (Fig. 3A), then declined as continued sea level rise submerged smaller islands below the 20 ha threshold. Total island area approached modern values early, around 8 ka, but continued to fluctuate until stabilising at 2ka (Fig. 3B). Indeed, the modern spatial configuration had largely stabilised by 3 ka (Fig. 2). When the size of the island-size threshold is varied, island number and area varied, but on the whole produced similar temporal dynamics (Fig. S4).

**Figure 3.**
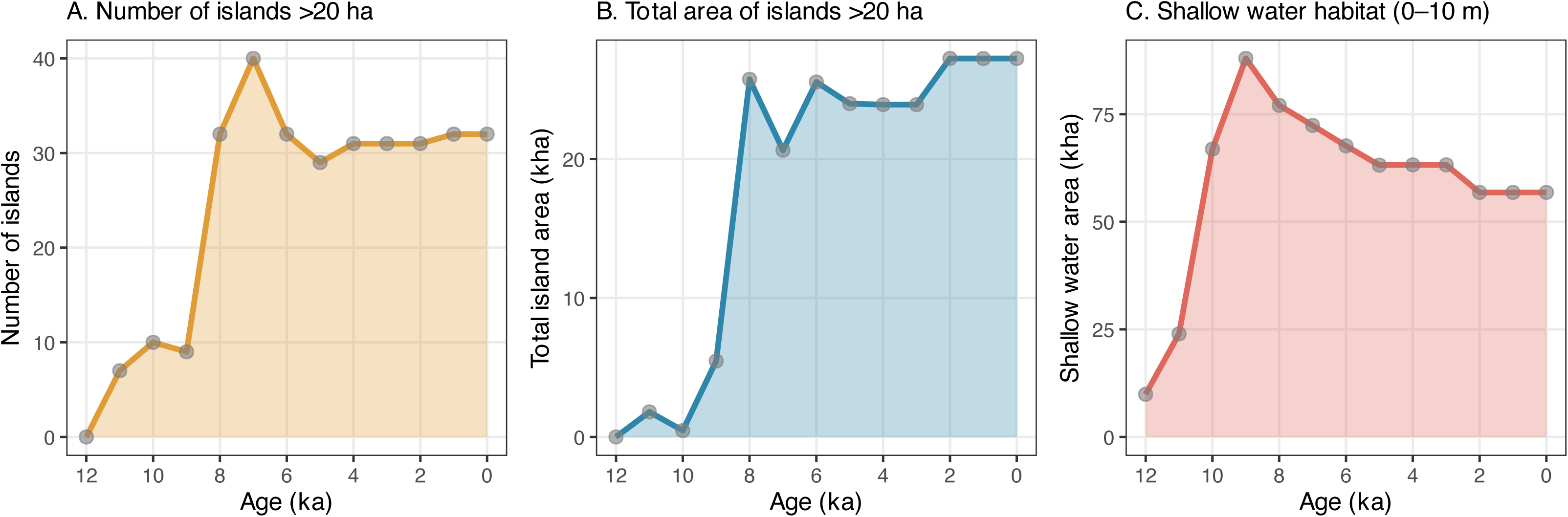
Estimated island and shallow water habitat dynamics reconstructed from post-glacial sea level rise over the past 12kyr in the Bocas del Toro archipelago, Panamá. (A) Total number of islands larger than 20 ha, and (B) Total land area of islands larger than 20 ha. The 20 ha threshold excludes digital elevation model (DEM) artefacts and reflects a biologically relevant island size. (C) Area of shallow water habitat (0–10 m depth) relative to contemporaneous sea level from 12 ka to the present, representing potential habitat for coral reefs, seagrass beds, and mangroves, i.e. three of the region’s most diverse and productive marine ecosystems.

Individual islands experienced markedly different isolation histories (Fig. 4, Table S1). The age of isolation from the mainland ranged from 9.5 ka (Escudo de Veraguas, the earliest) to 2.9 ka (Cristóbal, the most recent). Some islands revealed simple isolation trajectories such as early separation from the mainland followed by a sharp (e.g. Zapatillas) or more gradual (e.g. Escudo de Veraguas) area decline dependent on the bathymetric configuration surrounding the island. Other islands show more complex histories with multiple fragmentation events. Roldán and Pastores, for example, remained connected as a single landmass for nearly 7 kyrs after their separation from the mainland, and then isolated from each other into their modern configurations, producing a step-like decline rather than smooth exponential decay (Fig. 4).

**Figure 4.**
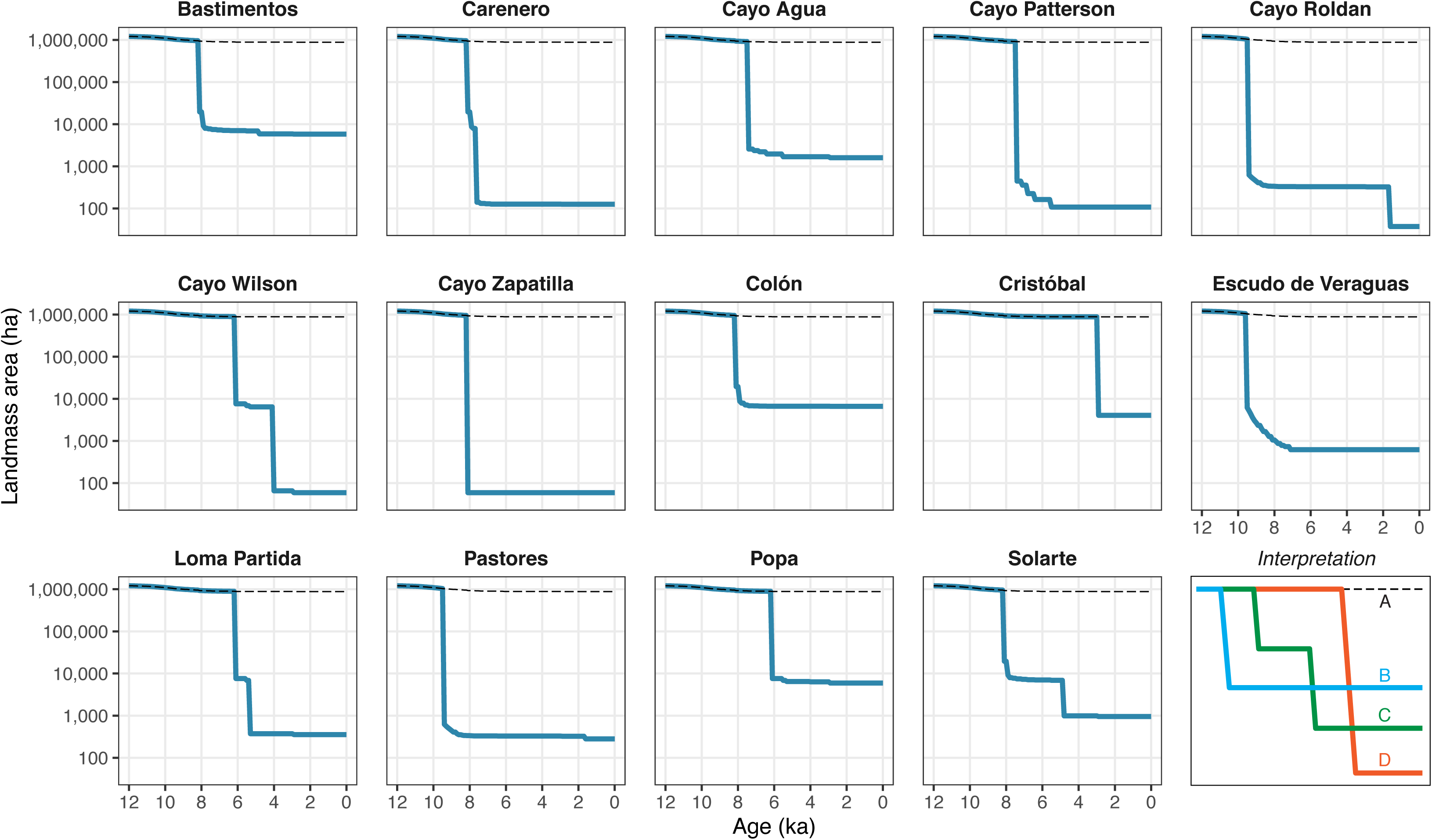
Temporal dynamics of island area during postglacial sea level rise in the Bocas del Toro Archipelago. Area trajectories for 14 islands reconstructed from bathymetric data and sea-level curves spanning the past 12 kyr. Each panel shows landmass area (ha) declining through time as rising seas fragmented the once-connected landscape. Horizontal dashed lines indicate mainland area. Abrupt drops in area represent when an island became isolated from either the mainland or a neighbouring landmass. Plateaus represent periods of stable island configuration. Interpretation panel (lower right) illustrates three archetypal trajectories: (A) mainland baseline; (B) early-isolated large island retaining substantial area (e.g., Bastimentos); (C) complex fragmentation history with initial early isolation, prolonged connection to a neighbouring island, then secondary isolation to medium size (e.g., Solarte); (D) late-isolated island (e.g., Cristóbal). These contrasting histories generate variation in isolation age, isolation area, and cumulative habitat availability (habitat-years), potentially useful context for biogeographic analyses.

Decay rates that quantify exponential area loss following isolation, varied considerably across islands, from near zero for Cayo Zapatilla to 1.3 per ka for Cayo Wilson (Table S1). Higher decay rates characterise islands either with shallow bathymetric profiles leading to prolonged losses over time, or groups of islands that are sequentially fragmented after their initial isolation from the mainland. The latter is also captured by the pairwise isolation matrix (Table S2) revealing that several island groups remained interconnected long after mainland separation. Roldán and Pastores isolated from the mainland at 9.4 ka but remained connected to each other until 1.6 ka, representing nearly 8 kyr of shared history post-mainland separation. Similarly, the island complex comprising Cayo Wilson, Loma Partida, and Popa were connected until 5.3–5.5 ka despite all three separating from the mainland by 6.1 ka.

### Shallow marine habitat Holocene dynamics

Our models reveal shallow water habitat (defined as 0-10m depth) followed a distinct temporal pattern from terrestrial habitat (Fig. 3C). At 12 ka, shallow habitat totalled only ∼10,000 ha, restricted to narrow strips along steep volcanic arc slopes lacking protected embayments. The absence of a drowned barrier reef in modern bathymetric data (Fig. S1) suggests these slopes were not conducive to barrier reef formation during the rapid early Holocene transgression.

Rapid expansion of shallow water habitat in the region began around 11 ka as rising seas flooded the gently sloping shelf (Figs. 3C). At 10 ka, major embayments including Laguna de Chiriquí had begun forming, creating extensive protected shallow platforms (Fig. 2). Shallow water habitat peaked at ∼88,000 ha around 9 ka, a nearly ninefold increase from early Holocene conditions achieved in less than 3,000 years. Since 9 ka, shallow water habitat declined progressively to ∼57,000 ha today, representing 65% of the mid-Holocene maximum (Table 2). This decline reflects both the drowning of outer shelf areas into depths beyond the 0–10 m range, as well as infilling of shallow bays through sediment accumulation and reef growth.

**Table 2.** Predicted habitat changes under 2150 sea level rise scenario. Comparison of current (2025) versus projected (2150) land area and shallow water habitat (0-10 m depth), with absolute and percentage changes. Scenario based on IPCC AR6 SSP3-7.0 projection combined with predicted tectonic subsidence (Plafker & Ward 1992).

| Scenario | Total land area (ha) | Total island area (ha) | Shallow water area (ha) |
| --- | --- | --- | --- |
| Current (2025) | 907,523 | 27,240 | 56,692 |
| Future (2150) | 864,803 | 24,070 | 85,211 |
| Change | -42,719 | -3,169 | 28,518 |
| Change (%) | -4.7 | -11.6 | +50.3 |

Terrestrial and marine trajectories were thus opposing through the Holocene (Fig. 3): marine habitat expanded rapidly then declined, whilst terrestrial habitat fragmented then stabilised. The extensive calm-water reef, seagrass and mangrove systems that characterise Bocas del Toro today are a product of currently high sea levels. Because marine organisms disperse effectively via pelagic larvae (Pinheiro et al. 2017), these shallow-water reconstructions quantify habitat availability rather than predict community structure.

### The Pleistocene context

The configurations of land and shallow water habitat we reconstructed above through the last 12 kyr represent just a brief interval in the much longer geological record of Bocas del Toro, and yet many biological processes unfold on such time scales of millions of years. Sea level reconstructions for the past one million years indicate a modal value of approximately −75 m relative to present, with the current interglacial highstand representing roughly 1% of that time span. The modern configuration of 32 islands exceeding 20 ha has therefore occurred rarely in the archipelago’s history (Fig. 5). For the majority of the time over the last 1 million years the region supported between zero and three islands > 20 ha (Fig. 5) and existed not as an archipelago but as a continuous coastal lowland extending from the mainland to what is now the outer shelf edge. And while the numbers of islands >20 ha had peaked at more than 80, sea level configurations with more than 25 islands occurred for less than 20,000 years (2%) in total over the last 1 Myr. In contrast, the six most frequently occurring sea level states, which together account for more than 31% of the past million years, all lack any substantial archipelago (Fig. S7).

**Figure 5.**
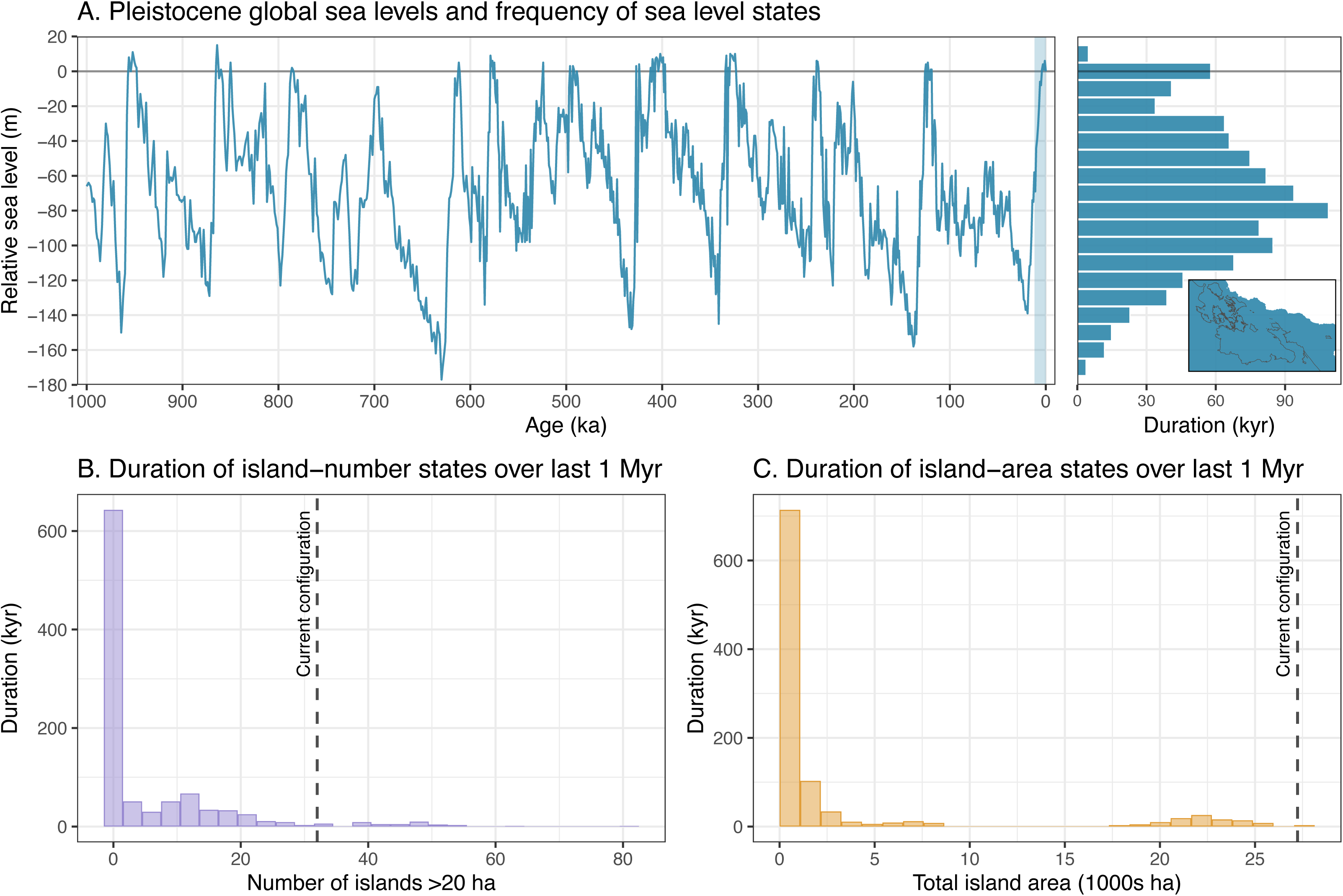
Pleistocene-Holocene island dynamics in Bocas del Toro, Panama (last 1 Myr). (A) Global sea level reconstruction based on Clark et al. (2025), with a marginal histogram showing the cumulative duration (kyr) spent at different sea level ranges (10 m bins). Green shading marks the Holocene (11.7–0 ka). The small inset map shows the land configuration of the Bocas del Toro region at the most frequently occurring sea level state over the last million years (5 m bins), illustrating that the most common historical condition lacked any substantial archipelago (see also Fig. S7). (B) Cumulative duration spent in different island-number states (islands >20 ha), calculated from digital elevation models at each sea level stand. (C) Cumulative duration spent in different total island-area states. Dashed lines in B and C indicate the modern configuration. The current archipelago has spent relatively little time in its present state; no islands at all was the most common historical condition.

These findings have archaeological implications. Humans arrived on the Isthmus between 25 and 16 ka (Ranere & Cooke 2021; Cybulski et al. 2025), and possibly earlier (Goebel et al. 2008), placing initial colonisation firmly within the Pleistocene. If early peoples were primarily coastal (Goebel et al. 2008), the majority of material evidence now lies submerged beneath modern sea level. Our models show they encountered a continuous lowland fronted by steep, high-energy shorelines (Fig. S7 map) under substantially drier conditions (Piperno et al. 1991a, 1991b; Piperno 2011). The protected lagoons, mangrove forests, and seagrass beds that define the archipelago today did not exist. The calm-water systems developed only during the mid- to late-Holocene transgression (Fig. 2). Stable human communities occupied these habitats from at least 4 ka until Spanish contact (Baldi 2011; Wake et al. 2013; Wake 2024), but the archipelago they inhabited clearly bore little resemblance to the Pleistocene coastline encountered by the region’s first peoples.

### Future habitat projections

Under a moderate emissions scenario (IPCC SSP3-7.0) combined with predicted tectonic subsidence, we project substantial but asymmetric changes to terrestrial and marine habitats by 2150 (Table 2). Our model predicts 12% loss of total island area across the archipelago, corroborating local flood modelling for Isla Colón that identified extensive inundation of low-lying coastal areas under similar scenarios (Ciniglio et al. 2021). These losses may be compounded by the intensifying flood risk recently predicted for the region (Petiangma et al. 2025).

This aggregate figure, however, obscures spatial inequality. The areas most vulnerable to inundation are those where human settlement concentrates. Bocas Town on Isla Colón sits almost entirely at sea level, and the narrow isthmus connecting the island’s northern and southern halves (carrying the main road) is projected to be severed under similar scenarios (Ciniglio et al. 2021). Elsewhere in Caribbean Panama, indigenous Guna communities on low-lying coral islands have already begun relocating to the mainland (Galindo Delgado & García Sánchez 2025).

In sharp contrast, shallow water habitat (0–10 m depth) is projected to expand by ∼50%, representing a return towards mid-Holocene levels (Table 2; Fig. 3C). Whether degraded Caribbean ecosystems can exploit this new space is doubtful because contemporary reefs are already failing to keep pace with current sea level rise (Perry et al. 2025), and thermal stress, disease, and declining water quality show no sign of abating. Rising seas thus simultaneously displace coastal communities and create marine habitat that degraded ecosystems may be unable to colonise.

### Biogeographical applications

To illustrate how the geomorphometric framework developed here can inform biological analyses, we examined species richness patterns of five terrestrial vertebrate groups across the archipelago using museum specimen records from USNM collections (see methods). The five groups (Anura, resident and non-migratory terrestrial birds, Chiroptera, Rodentia, and Squamata) span a broad range of dispersal capabilities: from frogs with limited capacity for sustained saltwater crossing (although see Measey et al. 2007), to bats that routinely traverse water tens of kilometres. Collecting effort varied substantially across islands and groups (Fig. S8), and sample sizes were small (7-12 islands), so we present these analyses as exploratory correlations rather than formal model comparisons. Nevertheless, the patterns that emerge are consistent, ecologically interpretable, and highlight both the strengths and limitations of different geomorphometric approaches in continental shelf archipelagos. Species richness varied substantially across the archipelago, with the largest islands (Colón, Bastimentos, Popa) supporting the most species across all groups, and several smaller islands and the highly isolated Escudo de Veraguas lacking sufficient records for most or all groups (Fig. 6A).

**Figure 6.**
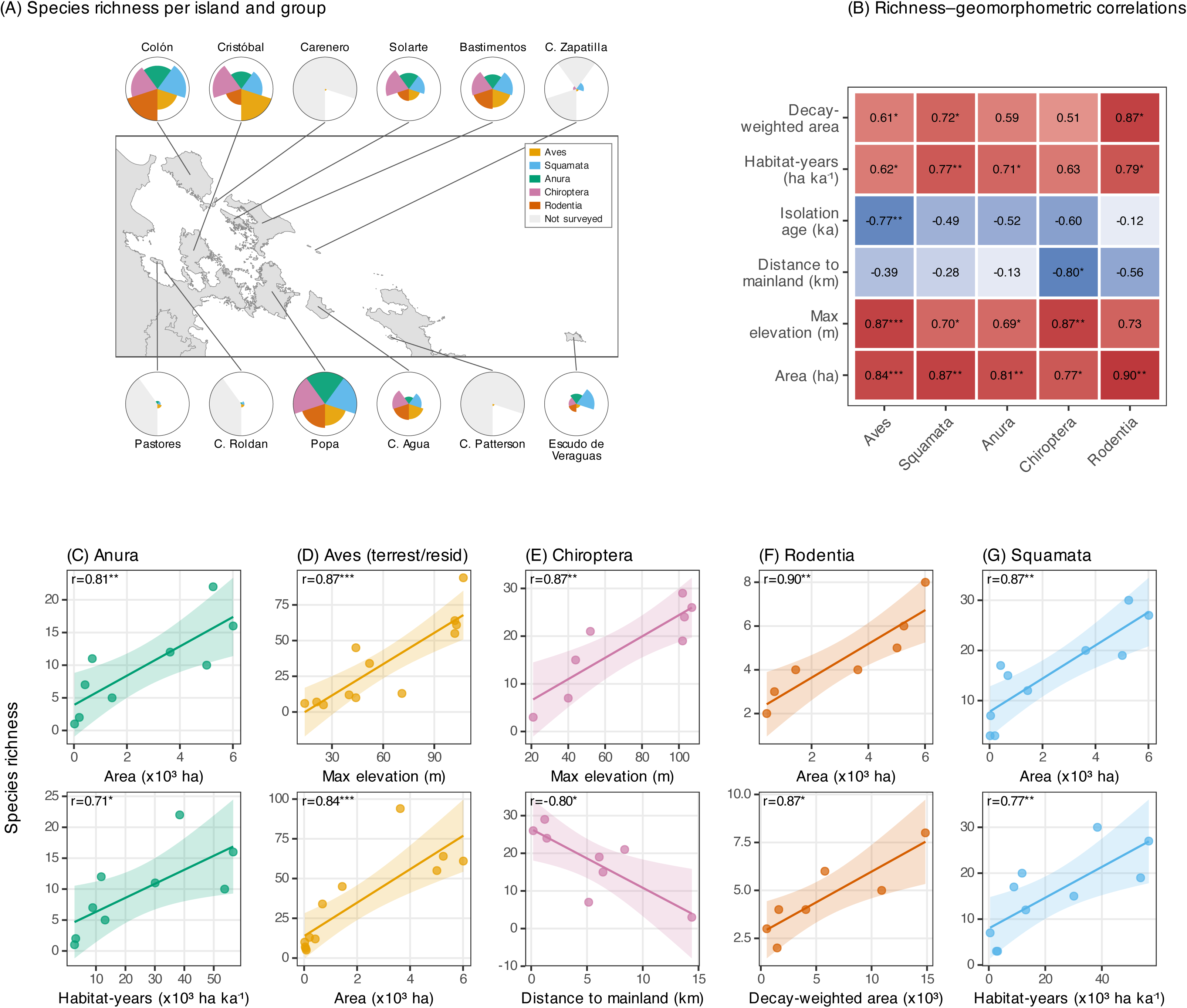
Species richness of terrestrial vertebrate groups across islands of the Bocas del Toro archipelago, Panamá. (A) Map of the archipelago showing species richness per taxonomic group for each island as pie charts (colours as inset key). *[Sector radii are log-transformed (log[x+1], scaled to the per-group maximum) to aid visual comparison across groups with very different richness ranges.]* A full-radius sector indicates the group maximum across all islands; groups absent from an island are shown as light grey segments. Lines connect each pie to its island centroid. (B) Heatmap showing Pearson correlation coefficients between species richness and six key geomorphometric variables for each taxonomic group. Values indicate correlation strength (r), with significance levels: *p < 0.05, **p < 0.01, ***p < 0.001. Colour intensity reflects correlation magnitude (red = positive, blue = negative). (C–G) Scatter plots showing the top two strongest correlations for each group: (C) Anura (frogs, n = 9 islands), (D) Aves (terrestrial and resident only, n = 12 islands), (E) Chiroptera (bats, n = 8 islands), (F) Rodentia (rodents, n = 7 islands), and (G) Squamata (reptiles, n = 10 islands). Each panel displays species richness against the predictor variable, with linear regression lines and 95% confidence intervals.

### Area and elevation

Present-day island area was the strongest single predictor of species richness across all five groups (Fig. 6B), consistent with the fundamental prediction of island biogeography theory. Maximum elevation performed nearly as well (significant for four of five groups, Figs. 6B, S9), likely reflecting the strong correlation between these two variables (larger islands tend to have higher elevation) and the role of topographic complexity in generating habitat heterogeneity. For Chiroptera, elevation may carry additional explanatory weight beyond its correlation with area: limestone karst, which provides roosting caves, occurs preferentially on higher-elevation islands (Coates et al. 2005). These results confirm that the classical species-area relationship holds robustly in this system, providing a baseline against which historical and composite metrics can be evaluated.

### Historical metrics extend power

Among the historical metrics, cumulative habitat availability since isolation (habitat-years) showed consistently strong positive correlations across all groups (Fig. 6B, S9). This metric integrates both how large an island has been and for how long, capturing the cumulative opportunity for populations to persist since mainland separation. Its broad predictive strength suggests that extinction dynamics in this system reflect not just current conditions but the entire post-isolation trajectory of habitat availability (Halley et al. 2016). Among the composite metrics, decay-weighted area, which discounts initial island area by the empirically derived rate of subsequent contraction, performed well for four of five groups and may be informative for taxa experiencing ongoing extinction debt, where current diversity reflects historical rather than present habitat configurations.

For resident terrestrial (non-migratory) birds, richness declined significantly with isolation age, meaning that islands separated from the mainland longest harbour fewer species. This negative relationship is weaker or absent in other groups (Figs. 6B, S9) and may potentially reflect progressive ecological relaxation (Valente et al. 2020) where bird communities captured at isolation gradually lose species as island areas contract, with the longest-isolated islands having experienced the greatest cumulative extinction. Alternatively, as our analysis excluded migratory species, older islands might support proportionally more migrants (which are not counted) competing for resources with residents during a portion of each year, effectively reducing the richness of the resident community we measure. The pattern could also reflect slow erosion of the mainland source pool available for recolonisation over the Holocene. Distinguishing among these hypotheses will require compositional analyses examining which species are lost from older islands and whether functional or phylogenetic diversity shows parallel declines. We note that with n = 12 islands in the bird analysis, statistical power to detect moderate correlations is limited, and the isolation age result should be treated as a hypothesis to test rather than a confirmed pattern.

### Isolation indices confounded by island size heterogeneity in this continental shelf setting

In contrast to the strong performance of area, elevation, and temporal metrics, several of the indices designed to quantify spatial isolation performed poorly or produced counterintuitive results. Distance to mainland was non-significant for four of five groups, achieving significance only for Chiroptera (r = −0.80, p = 0.018), the taxon with the greatest overwater dispersal capability. The proximity index, which weights surrounding island areas by inverse squared distance, was non-significant for all groups. Most strikingly, the buffer zone isolation indices (B1 through B64) showed weak, frequently negative correlations with richness across most groups (Fig. 6B), the opposite of the positive relationship expected if surrounding landmass facilitates colonisation.

We propose that the failure of these spatial metrics reflects a structural confound between island size and buffer values inherent to this archipelago’s geometry, rather than a fundamental limitation of the indices themselves. In Bocas del Toro, islands formed through progressive flooding of a continuous landscape, producing a nested spatial configuration where small islands are often remnant hilltops surrounded by (and close to) larger landmasses. As a result, small species-poor islands tend to have high buffer index values (because adjacent large islands fill their buffer zones), whilst large species-rich islands have low buffer index values (because their buffers extend primarily over open water). The result is a confound between island area and buffer isolation that inverts the expected biological signal. In oceanic volcanic archipelagos, where islands often form semi-independently and small islands surrounded by large neighbours are comparatively rare, area and surrounding landmass vary more independently and buffer indices perform as expected (Weigelt & Kreft 2013; Itescu et al. 2020). Researchers applying buffer-based or proximity-weighted isolation metrics to land-bridge, stepping-stone, or other continental shelf archipelagos should be alert to this confound.

### Dispersal capability structures taxon-specific responses

The contrasting responses across groups invite comparison of how dispersal biology mediates the influence of geomorphic history on diversity. Frogs (Anura), as obligate freshwater breeders with negligible saltwater tolerance (Galeano et al. 2023), effectively become stranded on islands at the moment of separation. Their diversity shows the strongest relationship with present-day area (r = 0.81) and essentially no relationship with isolation age (r = −0.51, non-significant), consistent with communities “frozen” by the initial fragmentation event and subsequently shaped primarily by area-dependent extinction (MacArthur & Wilson 2001; Sin et al. 2022). Bats, by contrast, show the strongest response to distance from mainland (r = −0.80) and elevation (r = 0.87), consistent with populations maintained by ongoing dispersal from the mainland and supported by habitat features (e.g. caves, forest complexity) correlated with island topography. For such vagile taxa, richness may reflect the balance between recurrent colonisation and local extinction, with island area setting an upper bound to richness and mainland distance modulating the supply of colonists. Rodents, squamates, and birds fall between these extremes in ways broadly consistent with their respective dispersal capabilities, though the small sample sizes and many limitations enforced by the nature of museum collection data preclude formal comparative tests at this point.

### Coda

These exploratory analyses demonstrate that the geomorphometric framework generates taxon-specific, testable hypotheses about how fragmentation history structures biodiversity. Teasing apart the relative contributions of correlated geomorphic variables will require modelling approaches (e.g. variance partitioning, phylogenetic comparative methods, or population genomic analyses) and more complete sampling, both beyond the scope of this paper. The metrics and reconstructions we provide supply the explanatory variables for such work, enabling researchers to move beyond simple species-area relationships towards process-based models incorporating temporal dynamics. For marine systems, the framework tracks when and where substrate became available for reef, seagrass, and mangrove colonisation, providing context for understanding community assembly, historical human exploitation of coastal resources, and the conservation of ecosystems whose current extent represents a geologically brief and unusual state. Extending biological analyses to marine specimen collections could enable analogous tests of how the dramatic expansion and subsequent contraction of shallow marine habitat documented here has structured the diversity and composition of shallow water reef and seagrass communities.

## Author contributions

Conceptualization: AO, MJB. Funding acquisition: AO, MJB, TJP, HLA. Data production: AO, MJB, CS, TJP. Investigation: All authors. Methodology: AO, SGAF. Code production: AO. Writing original draft: AO. Writing (review & editing): All authors.

## Supporting information

Supplementary information compiled

## Acknowledgements

Mr. Sebastian Castillo provided exceptional, professional field support and friendship to AO as his boat driver across the Bocas del Toro Archipelago for close to two decades. Javier Pardo, Kimberly García and many others supported the project. We thank Ximena Boza for constructive criticism of the manuscript. Chris Milensky, Addison Wynn and Michael McGowen provided USNM specimen data. Kevin de Queiroz updated species names for amphibians and reptiles. Charles O. Handley, Jr. and Storrs L. Olson (both now deceased) organized the Smithsonian Bocas Expeditions from 1987-1991 during which many of the specimen records utilized here were collected. We acknowledge their valuable contributions along with those of numerous field workers who participated in those expeditions. We thank Joyce and Mike Bytnar for their continued support of research on the Isthmus. Historical bathymetric data curation was provided by the Instituto Geográfico Nacional Tommy Guardia and the UK Hydrographic Office. We give special thanks to Milton Solano for guidance and support in geospatial analyses.

## Code and data availability

Complete R scripts and Python pipelines for all analyses are available on the GitHub and Zenodo repository (https://doi.org/10.5281/zenodo.18732423). The integrated DEM, island polygon shapefiles, sea level curves, calculated geomorphometric metrics, and shallow water habitat time series are provided to ensure full reproducibility. See Supplementary Methods for further details and links to external data sources.

