## Supplementary information compiled for "Geomorphic evolution of a Caribbean biological hotspot"

### Resumen (Spanish abstract)

Los archipiélagos de plataforma continental ofrecen oportunidades excepcionales para poner a prueba la biogeografía de islas en un contexto no equilibrado y con resolución temporal, porque el aumento postglacial del nivel del mar impulsa cambios fechables tanto en el área como en el aislamiento de las islas. Sin embargo, la mayoría de los sistemas costeros aún carecen de reconstrucciones que integren los cambios espaciales en la configuración insular con historias de fragmentación ponderadas por duración. Dicha información es esencial para interpretar los patrones genómicos y de biodiversidad modernos, y para evaluar respuestas a la fragmentación como la deuda de extinción, el retraso en la inmigración y las respuestas adaptativas a lo largo del tiempo. Aquí reconstruimos la historia geomórfica tierra-mar del Archipiélago de Bocas del Toro, Panamá, combinando batimetría y topografía de alta resolución con un modelo de nivel del mar corregido por sedimentación y movimiento tectónico. A partir de esto, cuantificamos el aislamiento a múltiples escalas, la dinámica histórica de fragmentación y métricas compuestas que integran espacio y tiempo. Mostramos que las islas actuales se aislaron secuencialmente entre 9,5 y 2,9 ka, con tasas de contracción de área posteriores al aislamiento que varían veinte veces entre islas. En ese mismo intervalo, el hábitat marino somero (0–10 m de profundidad) se expandió nueve veces a medida que la inundación de la plataforma creó espacio para arrecifes, pastos marinos y manglares. La extensión de los análisis al Pleistoceno revela que el archipiélago moderno es atípico y que durante gran parte del último millón de años la región existió como planicie costera continua en lugar de islas discretas. Las proyecciones bajo escenarios de emisiones moderadas predicen aproximadamente un 5% de pérdida de hábitat terrestre pero un 50% de expansión de hábitat marino somero para 2150, aunque es incierto si los arrecifes caribeños degradados pueden aprovechar esta expansión. Las correlaciones exploratorias entre la riqueza de especies de grupos de vertebrados terrestres y predictores geomorfométricos revelan que el área insular, la elevación máxima y la disponibilidad acumulativa de hábitat desde el aislamiento son los correlatos más sólidos de la diversidad, mientras que los índices de aislamiento por zona de amortiguamiento están confundidos por la covariación entre el tamaño de las islas y la masa terrestre circundante en este contexto de plataforma continental. En conjunto, Bocas del Toro constituye un experimento natural con calibración temporal para poner a prueba cómo la conectividad dinámica y el aislamiento han moldeado la biodiversidad insular y costera.

**Palabras clave:** Biogeografía insular, relaciones especie-área, archipiélago continental, fragmentación de hábitat, cambio del nivel del mar, paleogeografía

### Supplementary Methods

#### Sediment trap deployment and analysis

We quantified modern sediment accumulation rates to estimate historical channel depths throughout the Holocene reconstruction. Sediment traps were deployed at four reef sites across the Bocas del Toro archipelago: Gallinazo Point (Reef 1, Almirante Bay), Cristobal (Reef 2, Almirante Bay), Cristobal Village (Reef 3, Almirante Bay), and Cayo Agua (Reef 4, Chiriquí Lagoon). These sites were selected to capture variation in sedimentation regimes across the two main water bodies of the archipelago.

At each site, five replicate sediment traps were established at 2 m depth along a 20 m transect parallel to shore. Traps consisted of 25 cm long, 4 cm diameter PVC tubes that were capped at the base and hammered into the substrate, left protruding approximately 5 cm above the sea floor. This design minimises resuspension whilst allowing passive sedimentation to accumulate within the tube.

Traps were serviced every two months over the study period. At each collection, traps were carefully removed, capped, and replaced with new tubes. Sediment was returned to the laboratory, dried at 60°C to constant mass, and weighed. Sedimentation rates were initially calculated as g $m^{-2}$ $day^{-1}$ based on the trap's cross-sectional area and deployment duration.

To convert mass accumulation rates to linear accumulation rates (mm $year^{-1}$) required for our palaeo-bathymetric corrections, we used sediment density values for carbonate-siliciclastic mixtures characteristic of the Bocas del Toro region (O'Dea et al. 2007). The mean accumulation rate across all 20 traps was 0.46 ± 0.24 mm $year^{-1}$ (mean ± SD). This archipelago-wide mean was applied uniformly in our historical reconstructions to estimate sediment infilling of channels and shallow areas since the Last Glacial Maximum.

#### *In situ* channel depth measurements

The limited density of soundings in nautical charts meant that the narrowest channels between islands were not always adequately resolved by interpolation. In particular, several shallow, high-connectivity channels were spuriously infilled by the IDW interpolation, artificially connecting islands. To correct these artefacts, we manually edited six channels for which we had independent *in situ* depth measurements.

Channel depths were measured using three methods: (1) direct diving observations with depth gauges, (2) boat-based sonar transects, and (3) local ecological knowledge from experienced boat operators. Measurements represent the controlling depth (i.e. shallowest point or saddle) of each channel, as this determines the timing of island isolation during sea level rise.

The six corrected channels and their measured depths were:

1. Cayo Wilson–Isla Popa channel (1.2 m): Measured in situ directly with a dive computer in 2015
2. Western Loma Partida–Isla Popa channel (2.5 m): Measured with depth transect through channel in 2024

3. Eastern Loma Partida–Isla Popa channel (1.8 m): Estimated channel depth based on habitats in the saddle area and fact that boats can pass through with the motor down.
4. Isla Cristóbal–mainland channel (1.0 m): Measured as shallowest part of the saddle with depth transect through channel in 2024.
5. Solarte–Bastimentos channel (1.5 m): Measured from boat sonar in 2015.
6. Pastores–Roldán channel (0.5 m): Estimated following consultation with Captain Sebastián Castillo, an experienced local boat operator, who noted that the channel cannot be passed with an outboard motor lowered. Given that outboard motors typically sit at approximately 0.5 m depth, we assigned this as the maximum channel depth.

For each channel, we delineated a polygonal footprint from high-resolution satellite imagery and GPS waypoints, assigned a uniform depth equal to the measured controlling depth, and rasterised these polygons to the DEM grid. The resulting rasters were then overlaid on the original DEM so that depth values within each channel polygon were replaced by the field-based channel depth. This procedure ensured that key connections between islands remained submerged to their observed sill depths in the DEM, preventing spurious land bridges during Holocene sea level reconstructions.

#### Digital elevation model production

Previous geomorphic reconstructions of the archipelago (Summers et al. 1997) contained errors due to misreading the units on topographic charts, where depth soundings in fathoms were inadvertently interpreted as metres. This led to underestimates of the depths of channels between islands and as a result, inaccurate reconstructions of the timings of island isolations. In addition, previous reconstructions did not account for sediment channel infilling or vertical tectonic movements.

We created an integrated terrestrial–marine digital elevation model (DEM) by combining bathymetric and topographic data. Bathymetric data were digitized from 14 high-resolution nautical charts belonging to the US Army Map Service (AMS) Series E762, procured from the UK Hydrographic Office and the Instituto Geográfico Nacional Tommy Guardia of Panama. These charts, predominantly at a 1:50,000 and 1:25,000 scale, represent the most comprehensive historical surveys of the region prior to modern dredging.

Original soundings were recorded in both fathoms (on older charts) and meters (on newer charts). We manually digitised these as point features and standardized all values to meters below present Mean Sea Level (MSL). Charts were georeferenced in ESRI ArcGIS Pro using identifiable coastal features and bathymetric contours as control points. The original NAD27 (EPSG:26717) or WGS 1984 UTM Zone 17N (EPSG:32617) horizontal datums of each chart was reprojected to a unified WGS 1984 Web Mercator Auxiliary Sphere (EPSG:3857) projection. Between depth points, we created a continuous bathymetric surface using Inverse Distance Weighting (IDW) interpolation. To ensure seamless integration with terrestrial data, bathymetric rasters were resampled to match the 30-m cell size of the NASA Shuttle Radar Topography Mission 1 Arc-Second Global DEM (Jarvis et al. 2008). The merged topobathymetric DEM was projected to the WGS 1984 Web Mercator Auxiliary Sphere (EPSG:3857) to facilitate distance calculations and spatial analyses.

The final DEM has 30 m horizontal resolution and ~±1 m vertical uncertainty. The uncertainty reflects combined errors from chart accuracy specifications (we estimate typically ±0.5 m for modern surveys), georeferencing error (estimated ±0.3 m), and interpolation between soundings (estimated ±0.5 m). These vertical errors can propagate through our isolation age estimates, potentially shifting calculated isolation timing by several hundred years. However, without a higher-resolution DEM from dedicated multibeam surveys, these uncertainties cannot be reduced at this stage.

We then formulated a suite of spatial and temporal metrics that characterise key features of archipelago configuration and island isolation over time. These are classified as modern metrics (quantifying current configurations), historical metrics (quantifying temporal dynamics through time), and composite metrics (integrating spatial and temporal dimensions; Table 1).

#### Modern geomorphometrics

##### Island area

We calculated island area by first unifying all polygons belonging to each island using spatial union operations (to account for islands digitised across multiple map scales), then computing total area using the sf package (version 1.0-14) in R. Areas are reported in hectares.

##### Distance to mainland

We measured distance to mainland as the shortest straight-line (Euclidean) distance between each island and the nearest mainland coast, calculated using the `st_distance` function in sf (Fig. 1B,C). We defined mainland as any contiguous landmass exceeding 100,000 hectares, which in practice means the lower Central American Isthmus and excludes all islands.

##### Maximum elevation

Maximum elevation for each island was extracted by cropping the DEM to the island polygon (with a 100 m buffer to ensure complete coverage), masking to island boundaries, and identifying the maximum value. Elevations are reported in metres above mean sea level.

##### Proximity index

Following Kalmar & Currie (2006), we calculated a proximity index quantifying the potential for inter-island dispersal and gene flow. Their index summed the areas of all other islands within 64 kilometres, inversely weighted by squared distance. However, because the squared distance weighting creates extreme variation when applied to archipelagos with both very close (<10 m) and distant (>10,000 m) inter-island separations, we found it better to present the metric in log10 space:

$$\text{Proximity index} = \log10[\Sigma(Ai / di^2)]$$

where $A\mathrm{i}$ is the area in hectares of island $\mathrm{i}$ and $d\mathrm{i}$ is the shortest straight-line distance in metres from the target island to island i. All islands within 64 km are included in the summation. This log-transformation is applied during calculation, compressing the variation into a scale suitable for statistical analysis and comparison. Each unit in Proximity represents an order of magnitude difference in connectivity. Higher values indicate greater proximity to substantial landmasses that could serve as sources for colonisation or rescue effects.

#### Buffer zone isolation indices

We quantified multi-scale isolation following the cumulative buffer approach of Weigelt and Kreft (2013) and Itescu et al. (2020). For each island, we created circular buffers extending from the island margin at radii of 1, 4, 16, and 64 km. These distances follow a base-4 logarithmic progression designed to span the range of dispersal distances potentially relevant for the archipelago's biota. The maximum buffer of 64 kilometres was specifically chosen to ensure that even Escudo de Veraguas, the most isolated island (17.3 kilometres from nearest land), would have non-zero values in the largest buffer zone.

For each buffer zone, we calculated the proportion of the buffer area occupied by land (including both mainland and other islands, but excluding the target island itself):

$$\text{B index} = (\text{Area of land within buffer}) / (\text{Total buffer area})$$

Higher values indicate lower isolation (more surrounding land), whilst lower values indicate higher isolation. We calculated four cumulative buffer indices: B1 index (0–1 km), B4 index (0–4 km), B16 index (0–16 km), and B64 index (0–64 km).

This approach captures different spatial scales of connectivity (Table 1). The choice of buffer distances is necessarily somewhat arbitrary, as different taxa have different dispersal capabilities. However, this base-4 system provides consistent coverage across spatial scales appropriate for the Bocas del Toro system. Other systems should consider different buffer distances that are appropriate to the islands of choice, their source pools and the biological questions under consideration.

#### Historical geomorphometrics

We reconstructed changing island configurations throughout the Holocene (12 ka to present) by developing a sea level model and applying it to our terrestrial-marine integrated DEM at 100-year intervals .

##### Sea level curve reconstruction

We first developed a Caribbean-specific relative sea level (RSL) curve for the last 12 kyr following the methodology of Khan et al. (2017) and using Caribbean sea level data from Lambeck et al. (2014) and fitted a generalised additive model (GAM) with the form:

$$\text{RSL} \sim s(\text{Age}, k = 12, bs = \text{"tp"})$$

where RSL is relative sea level in metres, Age is years before present, $k = 12$ specifies the number of basis functions controlling curve smoothness, and $bs$ = "tp" indicates thin plate regression splines. We used restricted maximum likelihood (REML) estimation and applied a penalty parameter (gamma = 1.4) to favour smoother curves and reduce spurious oscillations.

The model was constrained to pass through 0 metres at present by including high-weight constraint points at 0, 1, and 2 ka (all with RSL = 0 metres and weights of 10,000, 500, and 100 respectively), ensuring the curve respects the fundamental definition that relative sea level equals zero at present day (Figure 3).

To quantify rates of sea level rise throughout the Holocene, we calculated mean rates for three time periods (12–8 ka, 8–4 ka, and 4–0 ka) following the temporal binning approach of Khan et al. (2017). For each period, we used the fitted GAM to predict RSL at the start and end points, then calculated the mean rate as the change in sea level divided by the duration of the interval, expressed as mm/year.

##### Sediment accumulation, uplift and subsidence corrections

Simply flooding the modern combined top-bathymetric DEM with past sea levels would fail to account for two key geological processes that alter palaeosurfaces: sedimentary infilling of channels and basins and net vertical tectonic movements from uplift and subsidence. Most historical sea level biogeographical studies neglect these processes despite their rates sometimes exceeding sea level rise itself (Khan et al. 2015, Govorcin et al. 2025). Our analyses aimed to incorporate these time-dependent corrections to more effectively reconstruct the palaeosurface existing at time $t$.

We estimated sediment accumulation rates using sediment trap data from twenty traps deployed at four shallow-water locations (Gallinazo Point, Cristobal, Cristobal Village, and Cayo Agua) with quarterly subsampling. Traps recorded deposition in g/area/day which we converted  to accumulation rates (mm/year) based on the density of the carbonate-siliciclastic sediments in the region (O'Dea et al. 2007). The mean rate across all traps was 0.46 (SD = 0.24) mm/year. This archipelago-wide mean was applied uniformly to estimate historical channel depths by subtracting accumulated sediment thickness from modern bathymetry, ignoring spatial variation in sediment rates.

The Bocas region experiences opposing tectonic forces: regional uplift from Cocos Ridge subduction (1.0 mm/year; Bennett et al. 2014) versus episodic localised subsidence from earthquake-related thrust faulting. Major earthquakes causing approximately 160 mm subsidence recur roughly every 170 years based on the 1991 Limón event and geological evidence (Plafker and Ward 1992, Suárez et al. 1995, Phillips and Bustin 1996), yielding a mean subsidence rate of 0.94 mm/year. The net vertical movement rate is therefore +0.06 mm/year (i.e. a slight net uplift).

This net vertical movement was added to the entire modern DEM before comparing to contemporaneous sea level. Sediment accumulation corrections were applied only to submerged areas within island footprints. Sediments fill channels over time, meaning they need to be removed to recreate past channel depth. Uplift and subsidence worked together to change entire island configurations and were also accounted for before attempting to recreate palaeosurfaces. Our DEM-based reconstructions do not account for terrestrial erosion. No erosion rate estimates are presently available for the low-lying terrain characteristic of the Bocas del Toro islands. Published denudation rates for the region (Gonzalez et al. 2016) have been derived from steep highland catchments where landslides dominate sediment flux and are therefore not directly applicable here. Locally derived erosion rates for low-relief terrain in the region would be required to incorporate this correction in future reconstructions.

#### Island area through time

At each time step, we identified emergent land by comparing the adjusted DEM to contemporaneous sea level. Cells with adjusted elevation above sea level were classified as land. We then used eight-directional connectivity analysis (where diagonally adjacent cells are considered connected) to identify spatially contiguous patches of land. For each modern island, we extracted all contiguous land patches that spatially intersected with that island's present-day polygon and summed their areas.

This approach automatically accounts for fragmentation and coalescence during sea level changes. When sea level was low, modern islands often formed parts of larger contiguous landmasses. As sea level rose, single landmasses then fragmented into multiple islands. Our method tracks these dynamics by identifying which modern islands belonged to which contiguous patches at each time step.

#### Isolation age and area

We determined when each island became isolated from the mainland by tracking the area of the contiguous landmass containing that island through time. Islands began as part of large coastal plains connected to the mainland. As sea level rose, some islands separated onto distinct landmasses whilst others remained connected to large areas.

Critically, we defined "mainland" dynamically at each time step as the single largest contiguous land patch in the region, acknowledging that the extent of mainland Panama varied through the Holocene as seas flooded the continental shelf. We defined isolation from the mainland as occurring when the landmass area dropped below 100,000 hectares representing clear separation from the isthmus and considerably larger than any island past or present.

We recorded both the timing of this isolation event (isolation age in thousands of years before present) and the area of the island at that moment (isolation area in hectares). We note that biological populations might functionally isolate before physical separation if intervening areas become inhospitable (e.g., flooded lowlands), or alternatively might maintain connectivity after physical separation through occasional over-water dispersal. Our metric provides the physical fragmentation baseline against which biological patterns can be compared.

##### Decay rate

Following isolation from the mainland, island areas continued to decline and separate as sea level rose. We quantified this contraction and fission by fitting an exponential decay model to the post-isolation area trajectory, following approaches used for other shelf archipelagos (Sin et al. 2022). For each island, we extracted area values from isolation time to 1 ka (after which time, no further changes in island connectivities occurred), every 1kyr. We then fitted a linear model to log-transformed areas:

$$\ln(\text{Area}) = \text{intercept} + k \times \text{time since isolation}$$

where $k$ is the decay rate parameter in hectares per thousand years. More negative values of $k$ indicate more rapid area loss. The resultant exponential model provides a simple one-parameter description of contraction dynamics. The decay rate parameter captures the average trend whilst allowing comparison across islands experiencing different geomorphic histories and captures some of the sequential isolations captured under the pairwise island isolation age approach (previously).

##### Habitat-years

We calculated cumulative habitat availability since isolation as the area under the curve of island area versus time, using trapezoidal integration across 100-year time steps. This metric, reported in hectare-thousands of years (ha·ka), integrates both the spatial dimension (how large was the island?) and temporal dimension (for how long?). It provides a single number summarising total habitat-time available since mainland disconnection and can be interpreted as the product of mean island area and time since isolation. This metric may be particularly relevant for understanding extinction debt (Halley et al. 2016), as species persistence depends not just on current area but on cumulative habitat availability through time.

#### Composite geomorphometrics

To integrate temporal and spatial dimensions of isolation history, we calculated three composite metrics combining isolation age and isolation area. These metrics may help test different hypotheses about how time and space jointly influence biological patterns.

##### Decay-weighted area

This metric incorporates the empirically derived decay rates to account for time-dependent area loss (Weigelt and Kreft 2013, Weigelt et al. 2016, Whittaker et al. 2017):

$$\text{Decay-weighted area} = \text{Isolation area} \times \exp(-k \times \text{Isolation age})$$

Where $k$ is decay rate (Table 1). This metric gauges "effective area" remaining after exponential decay. Islands with rapid decay rates (high $k$ values) and long isolation times receive greater temporal discounting than slowly contracting islands. It assumes that island area decline follows a predictable exponential trajectory, which we know is oversimplified but provides a first approximation. The metric is reported in hectares.

#### Log(time × area)

This metric uses an additive approach in log space from Weigelt et al. (2016) and Whittaker et al. (2017):

$$\text{Log(time} \times \text{area)} = \ln(\text{Isolation age [ka]} \times \text{Isolation area [ha]})$$

Note that this uses natural logarithm (ln). This formulation reflects the empirical finding that many island biogeographic relationships follow log-linear patterns, including species–area relationships and species–isolation curves. It weights time and area equally on a logarithmic scale, treating both as multiplicative factors contributing to habitat-time availability. The metric is dimensionless and provides a way to combine variables with different units.

#### Geometric mean

The geometric mean provides balanced integration of time and space:

$$\text{Geometric mean} = \sqrt{\text{Isolation age [ka]} \times \text{Isolation area [ha]}}$$

This treats temporal and spatial dimensions symmetrically, avoiding the dominance of either very large areas or very long isolation times that occur with simple multiplication. The geometric mean has units of √(hectare·ka) and provides an intermediate measure between the spatial and temporal extremes.

#### Biological interpretation and applications of composite metrics

These three composite metrics capture complementary aspects of isolation history and may be differentially predictive depending on the biological process of interest. Decay-weighted area emphasises recent habitat availability and may be most relevant for understanding contemporary demographic processes, extinction risk, and extinction debt, as it heavily discounts populations that experienced rapid area loss long ago. Log(time × area) treats temporal and spatial dimensions multiplicatively on a log scale, potentially making it most appropriate for evolutionary processes such as speciation and genetic divergence, where both extended isolation time and sufficient population size (captured by area) are necessary conditions. The geometric mean provides a neutral, symmetric integration of time and space without strong assumptions about their relative importance, making it useful for exploratory analyses when the relative contributions of time versus area are unclear. In practice, testing all three metrics against biological response variables (species richness, endemism, genetic diversity, population structure) can reveal which aspect of isolation history (e.g. recent effective area, multiplicative habitat-time availability, or balanced integration) best explains observed patterns for particular taxa or traits.

##### Shallow water habitat area through time

We estimated the area of potential habitat for coral reef, seagrass, and mangrove ecosystems through time by identifying seafloor at depths between 0 and 10 m below contemporaneous sea level. This depth range corresponds to the zone where these ecosystems most typically develop in the Caribbean (Briand et al. 2023). While reefs are known to develop at deeper depths, this range represents the zone where the majority of reef accretion occurs (Hynes et al. 2024). At each time step, we identified all DEM cells with adjusted elevations between sea level and 10 m below sea level, then summed their areas.

This represents an estimate of the potential maximum extent of shallow water habitat. Actual habitat extent depends on substrate type (coral reefs require hard substrate, mangroves require soft sediment, seagrasses require appropriate grain sizes), water quality, and other factors we did not model. However, the 0–10 m depth zone provides a consistent metric for tracking how shallow marine habitat availability changed through the Holocene.

##### Pairwise island isolation ages

To capture the complex sequential fragmentation history of the archipelago, we calculated pairwise isolation ages between all islands and between each island and the mainland. Whilst some islands isolated directly from the mainland and remained discrete entities, others separated from the mainland as part of larger landmasses that subsequently fragmented into multiple islands. This sequential isolation is biologically important because populations on islands that remained connected after mainland separation could continue exchanging individuals until their later separation from each other.

We determined pairwise isolation ages using connectivity analysis on the historical land masks. At each 100-year time step, we applied eight-directional patch analysis to identify spatially contiguous land areas. For each island pair, we tracked whether they shared patches (direct connection) or whether both remained connected to the mainland. Two islands were considered isolated from each other at the first time step when: (a) they no longer shared patches directly, and (b) at least one had lost connection to the mainland. This approach accounts for the fact that islands can exchange populations via mainland connections even when not directly adjacent. The resulting pairwise isolation matrix provides the age when each pair of islands became effectively isolated, capturing both direct separations and sequential fragmentations of initially larger landmasses.

#### Pleistocene context

To assess whether modern archipelago configurations are typical or unusual in a longer historical context, we extended our analyses to the last one million years using Pleistocene sea level reconstructions. We used the global mean sea level curve from Clark et al. (2025), which synthesises multiple proxy records and geophysical models to reconstruct sea level changes through the repeated glacial-interglacial cycles that characterised the Pleistocene. To visualise the frequency distribution of sea level states, we binned the Clark et al. (2025) sea level data into 10 m intervals and calculated the cumulative duration spent at each bin through the last million years. This approach was adapted from Flantua et al. (2019).

We then sampled 1001 time points through the last million years (every 1ka) from the Clark et al. (2025) dataset. For each time point, we applied the corresponding sea level to our uncorrected DEM and calculated the number of islands exceeding 20 hectares and their total area using the same methodology as our Holocene analysis: eight-directional connectivity analysis to identify land patches, with the largest patch designated as mainland and all other patches ≥20 ha classified as islands. This late Pleistocene -scale analysis provides lower resolution than the Holocene analysis, but offers a first-order approximation of archipelago dynamics through this time, acknowledging that actual configurations would have been modified by longer-term geological processes.

We generated frequency distributions showing how much time (in thousands of years) the archipelago spent in different configurations. For each state (number of islands or total island area), we calculated the duration by multiplying the number of sampled time points in that state by the mean interval between samples. This duration-weighting reveals which configurations were typical (persisted for long periods) versus unusual (brief or rare) through the Pleistocene.

This analysis does not capture every detail of island dynamics through the Pleistocene. The 1001 sampled time points miss most short-term fluctuations within glacial-interglacial cycles, and the lack of accretion corrections becomes increasingly problematic deeper in time. However, it provides a robust view of typical archipelago conditions and demonstrates how unusual the modern fragmented configuration is compared to the dominant Pleistocene states, when the region most commonly existed as continuous coastal lowland rather than as an archipelago.

#### Future sea level rise projections

To demonstrate the predictive utility of our geomorphic framework, we projected future habitat changes under a 2150 sea level rise scenario. We combined global sea level rise estimates from the IPCC Sixth Assessment Report SSP3-7.0 moderate emissions scenario (+1.34 m by 2150; IPCC 2021) with predicted tectonic subsidence specific to the Bocas region.

The 1991 Limón earthquake caused approximately 900 mm subsidence in the Bocas mainland, and an average of 150 mm in the archipelago (Plafker and Ward 1992), with a similar event in the early 19th century suggesting a recurrence interval of approximately 170 years (Suárez et al. 1995). Given this pattern and our 125-year projection period (2025–2150), we incorporated an additional 0.30 m subsidence (medium probability of two similar earthquakes based on the ~170-year recurrence interval), yielding a total relative sea level rise of +1.64 m by 2150.

We applied this future sea level to our present-day adjusted DEM using the same methodology as our historical reconstructions. We calculated the total land area by identifying all cells with elevations above projected sea level. Shallow water habitat (0–10 m depth) was calculated by identifying cells between projected sea level and 10 m below it. These projections assume no additional sediment or coral accretion over the next 125 years, given ongoing coral reef decline throughout the Caribbean and demonstrated inability of contemporary Caribbean reefs to keep pace with projected sea level rise rates (Perry et al. 2025).

### Museum specimen collation and standardisation

#### Data sources

Specimen records were drawn from three collections held at the US National Museum of Natural History (USNM), Smithsonian Institution: mammals (n = 4,492), herpetofauna (n = 3,001), and birds (n = 3,899). Records were collated and processed through a pipeline comprising island assignment, taxonomic classification, identifier extraction, metadata parsing, locality reclassification, and expert-recommended modifications. Following these procedures, the final dataset comprised 11,378 specimens (4,491 mammals, 3,000 herps, 3,887 birds) representing 624 unique species-level taxa. The pipeline is implemented in Python and available in our GitHub repository.

#### Island assignment

Each specimen was assigned to an island based on its raw locality string using a hierarchy of regular expression patterns. Fourteen canonical island names were defined, each with associated aliases, common misspellings, and English/Spanish variants. Patterns were applied in sequence, with the final match taking precedence; this allowed more specific patterns (e.g. Cayo Zapatilla) to override broader ones (e.g. Bastimentos). Source datasets required different field mappings: for mammals, island was inferred primarily from the Precise Locality field; for herpetofauna, from Locality; and for birds, from Precise Location, with fallback to county-level fields. Localities containing "Between Isla..." were flagged as ambiguous.

#### Taxonomic classification

Class and Order were assigned using source-specific approaches. For mammals, Order was taken directly from the source data, with one correction applied (*Metachirus myosuros*, reassigned to Order Didelphimorphia). For herpetofauna, taxonomy was resolved first against a curated species reference sheet, then via a genus-level lookup table encompassing over 60 genera; identifications to family level only (e.g. centrolenid, dendrobatid) were assigned Class and Order accordingly. For birds, Order was derived from Family via a lookup table of 54 families following AOU classification.

Binomial names were extracted from the taxon field using a set of explicit rules. Trinomials were reduced to species level; bracketed synonyms and author citations were removed. Two cases required specific treatment: specimens recorded as *Chalybura urochrysia isaurae × melanorrhoa* were collapsed to species level, as Bocas del Toro is the type locality for this subspecific hybrid zone; specimens of Manacus vitellinus × candei were retained as hybrids, reflecting over three decades of taxonomic research on this interspecific cross centred on the archipelago. Identifications to family level only were retained without binomial assignment (n = 40).

#### Specimen identifiers and collection metadata

USNM catalog numbers were extracted from source-specific column names and retained in positional order. Collector names were taken directly from source fields. Collection dates were parsed to ISO 8601 format at the precision available in each record (full date, month-year, or year). Parenthetical annotations and time information were stripped prior to parsing; for date ranges, the start date was retained. Dates that could not be parsed were recorded as null (n = 50).

##### Locality reclassification

Following initial island assignment, a second-pass reclassification corrected assignments where locality strings contained sufficient geographic detail to override the pattern-based result. Key corrections included the reassignment of 232 specimens across 14 localities to Colón (including Boca del Drago and eastern-coast points previously assigned to "Other"), a typo correction reassigning one locality to Bastimentos, and a spelling-variant correction reassigning one locality to Pastores. One locality ("Quebrada Pastores, on coast W of Isla Pastores") was reclassified to “mainland” based on its explicit geographic description. Two additional island categories were identified: Cayo Garcia (7 specimens across 3 localities; a distinct cay in Chiriqui Lagoon) and Tiger Cays (21 specimens across 5 localities; offshore rocks south of Peninsula Valiente). Fourteen localities (44 specimens) were flagged as ambiguous, with notes documenting the geographic basis for uncertainty.

##### Expert-recommended modifications

Following review of the collated dataset by taxonomic specialists (M. Braun, T. Parsons, K. de Queiroz), four categories of records were modified. First, 14 specimens were excluded: one genus-level identification that would artificially inflate island richness (*Glossophaga* sp. from Popa), one holotype with uncertain locality (*Sibon longifrenis*, 1909), five swifts with impossible collection logistics (same collector, date, but multiple distant localities), one misidentified specimen whose voucher has been lost (*Habia rubica*), and six water birds inappropriate for terrestrial island fauna analyses (*Larus sp., Stercorarius parasiticus*). Second, 454 *Manacus* specimens across multiple species and hybrids were collapsed to the single taxon "Manacus sp.", as genetic data demonstrate that all local populations are panmictic or nearly so and thus represent a single evolutionary unit for diversity analyses (ref needed). Third, 216 specimens from "Boca del Drago" were flagged as ambiguous unless the locality explicitly mentioned "Isla Colón" or "island"; this toponym refers both to a village on Colón and to the strait/channel between the island and mainland, creating positional uncertainty. Fourth, 27 specimens identified as *Lepidoblepharis* sp. or *L. buchwaldi* were consolidated to *L. xanthostigma*, as examination of field notes and specimens confirmed all records represent this species. These modifications reduced the final dataset to 11,378 specimens representing 624 unique binomials.

#### Biogeographic analyses and species richness map

Species richness per island was calculated for each of five terrestrial vertebrate groups (Anura, Aves, Chiroptera, Rodentia, Squamata) from the collated museum specimen dataset. Only island-group combinations represented by five or more specimens were included, to reduce the influence of incidental or survey-incomplete records on richness estimates. Species identity was determined from the standardised binomial names in the collated dataset.

To visualise spatial variation in species richness across the archipelago (Fig. 6A), we constructed a proportional pie-chart map in R using ggplot2 and the scatterpie package. For each island, a pie chart was positioned at the island centroid. Sector radii scaled to the per-group maximum richness across all islands, so that a full-radius sector indicates the highest observed richness for that taxonomic group. Islands with no specimens recorded for a group are shown as light-grey segments. Lines connect each pie chart to its island centroid. The colour scheme is consistent with the correlation coefficient plots (Figs. 6C-G).

Pearson correlation coefficients were calculated between species richness and six key geomorphometric variables (island area, distance to mainland, isolation age, isolation area, habitat-years, and the log[time × area] composite metric) for each taxonomic group. Significance was assessed at $p < 0.05$, 0.01, and 0.001 thresholds. Results are displayed as a heatmap (Fig. 6B) and as scatter plots with linear regression lines and 95% confidence intervals for the two strongest predictors per group (Fig. 6C–G).

### Code and data

To ensure the reproducibility of the analyses, tables, and figures presented in this study, all necessary data and code are available on Zenodo (https://doi.org/10.5281/zenodo.18732423).

#### External data

- Integrated terrestrial-marine DEM (30 m resolution) available from (Titcomb and O'Dea 2020).
- Caribbean sea level data from Lambeck et al. (2014).
- Pleistocene sea level reconstructions from (Clark et al. 2025).
- SRTM 30 m DEM from (Jarvis et al. 2008)
- Bathymetric data digitised from UK Hydrographic Office and Instituto Geográfico Nacional Tommy Guardia nautical charts (AMS Series E762)
- Sediment accumulation rates (C. Schloeder)
- Tectonic parameters from (Plafker and Ward 1992, Suárez et al. 1995, Bennett et al. 2014).

#### R Scripts

All analyses were conducted in R version 4.3.1 (or later) using the following packages: sf version 1.0-14 (spatial data handling), terra version 1.7-46 (raster processing), mgcv version 1.9-0 (GAM modelling), dplyr version 1.1.3 and tidyr version 1.3.0 (data manipulation), readr version 2.1.4 (data input), units version 0.8-4 (unit conversions), ggplot2 version 3.4.4, patchwork version 1.1.3, and tidyterra version 0.5.2 (visualisation), GGally version 2.1.2 and ggrepel version 0.9.4 (extended plotting), furrr and future (parallel processing).

Analysis scripts should be run in the following order:

1. 01_holocene_analysis.R – Reconstructs Holocene terrestrial dynamics including sea level modelling, sediment and tectonic corrections. Produces historical land masks at 100-year intervals and island-level metrics.

2. 03_modern_geomorphometrics.R – Calculates contemporary island geomorphometrics (area, distance to mainland, elevation, proximity index, buffer zone isolation indices).

3. 02_holocene_figures.R – Generates all Holocene figure outputs (Figs 2–4), including the combined bathymetric/palaeogeographic reconstruction, island area metrics through time, and island decay plots.

4. 04_island_iso_matrix.R – Calculates pairwise isolation ages between all islands (depends on land masks from script 01)

5. 05_pleist_multicore.R – Extends analysis to the last 1 million years using Clark et al. (2025) sea level reconstructions (note, uses parallel processing).

6. 06_future.R – Projects future habitat changes under 2150 sea level rise scenario (depends on outputs from scripts 01 and 03)

7. 07_bio_analysis.R – Filters taxonomic groups from USNM museum collections and correlates species richness with geomorphometric variables. Produces Fig. 6.

8. 08_tables.R – Combines modern and historical metrics, calculates composite geomorphometrics, generates Tables 1 and 2 in the manuscript.

9. 09_supplements.R – Generates supplementary analyses and figures

#### Python pipeline

Museum specimen data were collated from three separate USNM source files (mammals, herpetofauna, birds) into a single standardised dataset using a reproducible Python pipeline (00_collation_pipeline.py). The pipeline processes raw catalogue exports and produces two output files: a collated specimen table (bocas_NMNH_records_collated.csv; 11,392 specimens) and a locality-to-island mapping (bocas_localities_island_mapping.csv).

Island assignment uses regex pattern matching against 14 canonical island names applied to raw locality strings, with a "last match wins" specificity rule (e.g. "Bastimentos Island, Little Zapatilla Key" correctly resolves to Cayo Zapatilla rather than Bastimentos). A second pass applies manual reclassifications for localities that the regex could not resolve unambiguously; each correction is documented with geographic rationale. Localities that remain ambiguous (e.g. "Boca del Drago", which refers to both a village on Isla Colon and the adjacent strait) are flagged.

Taxonomy (Class, Order) is sourced per dataset: mammals from the USNM source "Order" column; herpetofauna from a tidy reference sheet with genus-level fallback; birds via AOU Family-to-Order mapping. Binomial names are extracted using rule-based parsing that handles family-level identifications (returned as null), genus-level records (retained as "Genus sp."), subspecies (reduced to binomial), author citations (stripped), synonym genera in brackets (removed), and explicit hybrid/intergrade cases. Collecting dates are parsed to the highest available ISO resolution (YYYY-MM-DD, YYYY-MM, or YYYY).

Expert-recommended modifications include: removal of problematic records (water birds inappropriate for terrestrial island fauna analysis, specimens with unresolvable locality or identification issues); collapsing all

Manacus taxa to a single operational unit (genetic data confirm local populations are panmictic); and consolidation of Lepidoblepharis sp. and L. buchwaldi to L. xanthostigma based on specimen re-examination.

The pipeline requires Python 3.8+ with pandas, openpyxl, and xlrd.

#### Derived datasets

The analysis scripts produce several derived datasets archived in the repository. Five species-by-island abundance matrices (one per taxonomic group: Anura, Chiroptera, non-migratory terrestrial Aves, Rodentia, Squamata) record specimen counts for each species on each of the 14 focal islands, along with an accompanying summary table (abundance_matrices_summary.csv). A tidy specimen-level dataset (bocas_species_tidy.csv) contains all filtered records used in the biogeographic analyses. Pearson correlation coefficients between species richness and all 15 geomorphometric predictors are recorded in bocas_correlations_richness_geomorph.csv. Island-group combinations with fewer than five specimens were excluded from correlation analyses.

### Supplementary tables

Table S1. Modern, historical and composite island geomorphometrics. Modern day geomorphometric characteristics and isolation metrics for islands in Bocas del Toro archipelago, Panama. Island metrics include area, distance to nearest mainland, maximum elevation, proximity index (inverse distance-weighted sum of nearby island areas; following Kalmar and Currie (2006), and cumulative buffer isolation indices at four spatial scales (B1 = 0-1 km, B4 = 0-4 km, B16 = 0-16 km, B64 = 0-64 km buffer zones; following Weigelt and Kreft (2013)), which represent the proportion of land within each buffer zone, where higher values indicate lower isolation. Proximity index calculated for islands within 64 km only. Explanation of all metrics and a short description of their calculation is given in Table 1. See Methodology for full descriptions.

| | Modern geomorphometrics | | | | | | | | Historical geomorphometrics | | | | Composite geomorphometrics | | |
|---|---|---|---|---|---|---|---|---|---|---|---|---|---|---|---|
| Island | Area (ha) | Distance to mainland (km) | Maximum elevation (m) | Proximity index | B1 index | B4 index | B16 index | B64 index | Isolation age (ka) | Isolation area (ha) | Decay rate (k) | Habitat-years (ha ka-1) | Decay-weighted area | Log(Time x Area) | Geometric mean |
| Bastimentos | 5012.5 | 6.2 | 102 | -1.7 | 0.02 | 0.07 | 0.15 | 0.35 | 8.1 | 19526.0 | 0.0719 | 53701.00 | 10906.5 | 12.0 | 397.7 |
| Carenero | 80.9 | 12.6 | 25 | -0.67 | 0.06 | 0.23 | 0.22 | 0.37 | 8.1 | 19526.0 | 0.2483 | 6356.00 | 2613.0 | 12.0 | 397.7 |
| Cayo Agua | 1439.7 | 6.5 | 44 | -3.2 | 0.00 | 0.01 | 0.13 | 0.40 | 7.4 | 2563.7 | 0.0622 | 13081.00 | 1618.0 | 9.9 | 137.7 |
| Cayo Patterson | 60.7 | 0.3 | 14 | -4.46 | 0.11 | 0.38 | 0.18 | 0.39 | 7.4 | 445.0 | 0.1619 | 1053.00 | 134.3 | 8.1 | 57.4 |
| Cayo Roldan | 22.3 | 1.3 | 44 | -1.66 | 0.02 | 0.28 | 0.58 | 0.44 | 9.4 | 623.6 | 0.1413 | 2716.00 | 165.2 | 8.7 | 76.6 |
| Cayo Wilson | 57.5 | 2.0 | 23 | 1.79 | 0.29 | 0.47 | 0.27 | 0.41 | 6.1 | 7596.1 | 1.3237 | 14199.00 | 2.4 | 10.7 | 215.3 |
| Cayo Zapatilla | 40.5 | 14.4 | 21 | -3.27 | 0.00 | 0.00 | 0.14 | 0.35 | 8.1 | 59.2 | 0.0000 | 480.00 | 59.2 | 6.2 | 21.9 |
| Colón | 6012.0 | 1.5 | 103 | -2.41 | 0.01 | 0.04 | 0.21 | 0.36 | 8.1 | 19526.0 | 0.0338 | 56588.00 | 14849.5 | 12.0 | 397.7 |
| Cristóbal | 3631.2 | 0.4 | 107 | -3.25 | 0.03 | 0.10 | 0.45 | 0.41 | 2.9 | 4063.2 | 0.0000 | 11783.00 | 4063.2 | 9.4 | 108.6 |
| Escudo de Veraguas | 419.0 | 17.3 | 40 | -5.41 | 0.00 | 0.00 | 0.00 | 0.27 | 9.5 | 6182.0 | 0.1511 | 8889.00 | 1471.4 | 11.0 | 242.3 |
| Loma Partida | 341.5 | 0.1 | 116 | -0.24 | 0.19 | 0.31 | 0.27 | 0.44 | 6.1 | 7596.1 | 0.4626 | 7498.00 | 451.9 | 10.7 | 215.3 |
| Pastores | 201.7 | 1.1 | 71 | -2.57 | 0.01 | 0.21 | 0.60 | 0.44 | 9.4 | 623.6 | 0.0334 | 3119.00 | 455.6 | 8.7 | 76.6 |
| Popa | 5254.7 | 1.2 | 102 | -0.15 | 0.05 | 0.10 | 0.16 | 0.41 | 6.1 | 7596.1 | 0.0448 | 38427.00 | 5779.7 | 10.7 | 215.3 |
| Solarte | 692.0 | 9.0 | 52 | -0.93 | 0.07 | 0.26 | 0.23 | 0.37 | 8.1 | 19526.0 | 0.4448 | 30098.00 | 532.0 | 12.0 | 397.7 |

Table S2. Pairwise island isolation ages in the Bocas del Toro Archipelago. Matrix showing the age (years BP) when each pair of islands became isolated from each other, calculated from connectivity analysis of historical land masks incorporating sea level rise, sediment accumulation, and tectonic corrections. Values represent the timing when island pairs could no longer exchange populations either through direct land connection or via shared mainland connectivity. Note that some islands show substantially different mainland isolation ages (first column) versus pairwise isolation ages, revealing sequential fragmentation history. See Methods for full calculation details.

| | **Mainland** | **Escudo de Veraguas** | **Cayo Roldan** | **Pastores** | **Bastimentos** | **Carenero** | **Cayo Zapatilla** | **Colón** | **Solarte** | **Cayo Agua** | **Cayo Patterson** | **Cayo Wilson** | **Loma Partida** | **Popa** | **Cristóbal** |
|---|---|---|---|---|---|---|---|---|---|---|---|---|---|---|---|
| **Mainland** | 0 | 9500 | 9400 | 9400 | 8100 | 8100 | 8100 | 8100 | 8100 | 7400 | 7400 | 6100 | 6100 | 6100 | 2900 |
| **Escudo de Veraguas** | 9500 | 0 | 9500 | 9500 | 9500 | 9500 | 9500 | 9500 | 9500 | 9500 | 9500 | 9500 | 9500 | 9500 | 9500 |
| **Cayo Roldan** | 9400 | 9500 | 0 | 1600 | 9400 | 9400 | 9400 | 9400 | 9400 | 9400 | 9400 | 9400 | 9400 | 9400 | 9400 |
| **Pastores** | 9400 | 9500 | 1600 | 0 | 9400 | 9400 | 9400 | 9400 | 9400 | 9400 | 9400 | 9400 | 9400 | 9400 | 9400 |
| **Bastimentos** | 8100 | 9500 | 9400 | 9400 | 0 | 7900 | 8100 | 7900 | 4800 | 8100 | 8100 | 8100 | 8100 | 8100 | 8100 |
| **Carenero** | 8100 | 9500 | 9400 | 9400 | 7900 | 0 | 8100 | 7600 | 7900 | 8100 | 8100 | 8100 | 8100 | 8100 | 8100 |
| **Cayo Zapatilla** | 8100 | 9500 | 9400 | 9400 | 8100 | 8100 | 0 | 8100 | 8100 | 8100 | 8100 | 8100 | 8100 | 8100 | 8100 |
| **Colón** | 8100 | 9500 | 9400 | 9400 | 7900 | 7600 | 8100 | 0 | 7900 | 8100 | 8100 | 8100 | 8100 | 8100 | 8100 |
| **Solarte** | 8100 | 9500 | 9400 | 9400 | 4800 | 7900 | 8100 | 7900 | 0 | 8100 | 8100 | 8100 | 8100 | 8100 | 8100 |
| **Cayo Agua** | 7400 | 9500 | 9400 | 9400 | 8100 | 8100 | 8100 | 8100 | 8100 | 0 | 7400 | 7400 | 7400 | 7400 | 7400 |
| **Cayo Patterson** | 7400 | 9500 | 9400 | 9400 | 8100 | 8100 | 8100 | 8100 | 8100 | 7400 | 0 | 7400 | 7400 | 7400 | 7400 |
| **Cayo Wilson** | 6100 | 9500 | 9400 | 9400 | 8100 | 8100 | 8100 | 8100 | 8100 | 7400 | 7400 | 0 | 5500 | 5300 | 6100 |
| **Loma Partida** | 6100 | 9500 | 9400 | 9400 | 8100 | 8100 | 8100 | 8100 | 8100 | 7400 | 7400 | 5500 | 0 | 5500 | 6100 |
| **Popa** | 6100 | 9500 | 9400 | 9400 | 8100 | 8100 | 8100 | 8100 | 8100 | 7400 | 7400 | 5300 | 5500 | 0 | 6100 |
| **Cristóbal** | 2900 | 9500 | 9400 | 9400 | 8100 | 8100 | 8100 | 8100 | 8100 | 7400 | 7400 | 6100 | 6100 | 6100 | 0 |

### Supplementary Figures

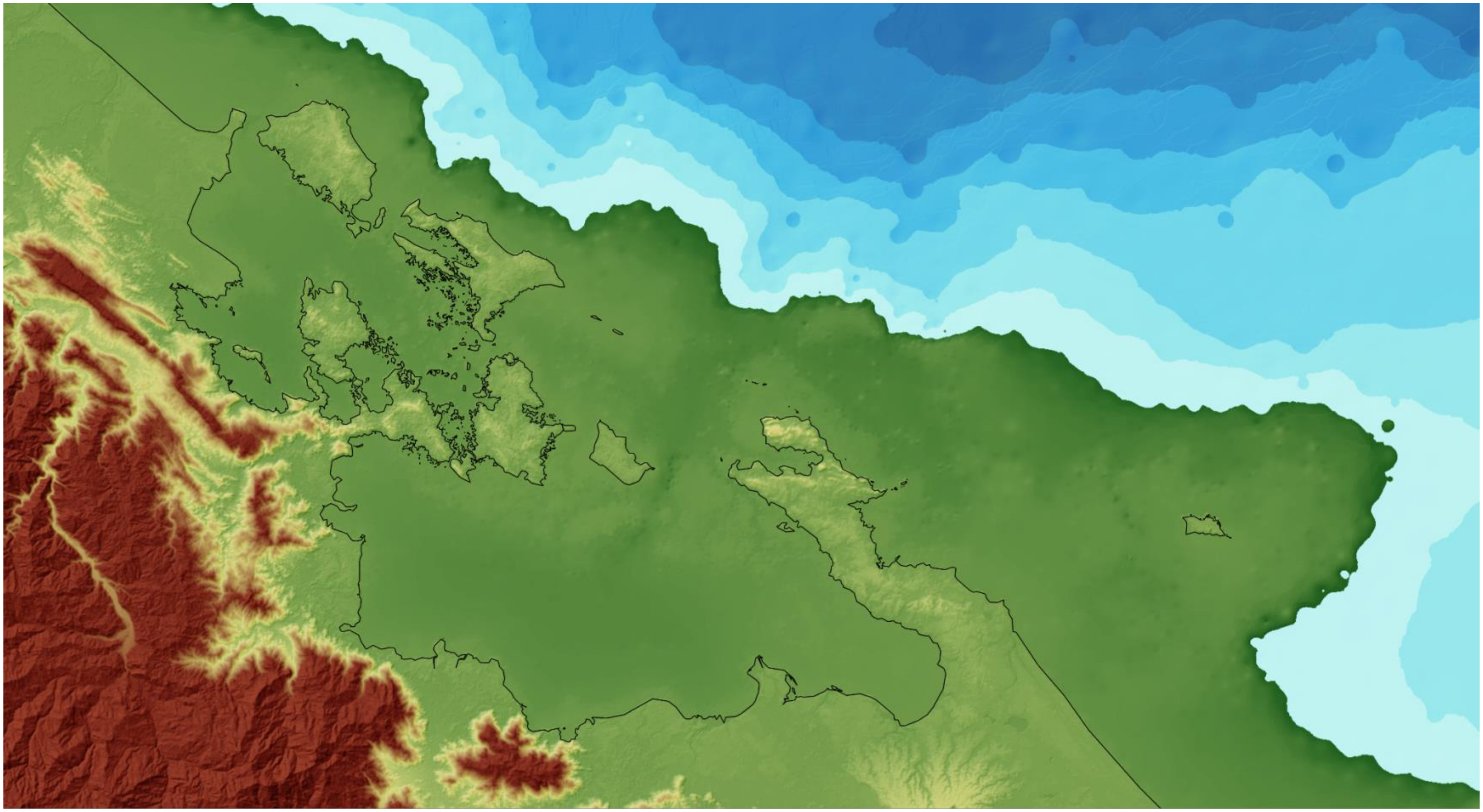

**Fig. S1. Bocas at the Last Glacial Maximum with sea level 125m lower than today.** Thin black line indicates modern day coastline.

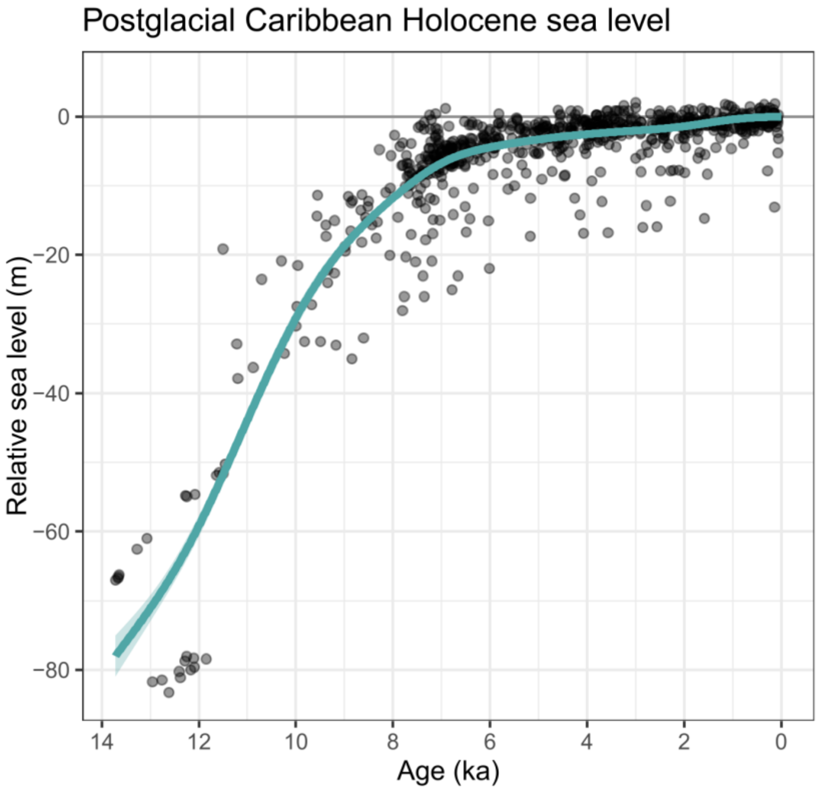

**Figure S2. Caribbean sea level reconstruction since 14ka.** Relative sea level (RSL) data from the Caribbean (Lambeck et al. 2014). The solid cyan line represents a generalised additive model (GAM) fitted to compaction-corrected data with ±1 standard error band (shaded area). The model is constrained to pass through 0 m at present (0 ka) and uses temporal smoothing to capture sea level changes from the late Pleistocene (pre 11.7 ka) through the Holocene (post 11.7 ka). Data span the period of postglacial marine transgression when rising sea levels reshaped the Bocas del Toro archipelago geography and island connectivity (Fig. 4). See Methodology and Approach and Supplemental Methods for full details.

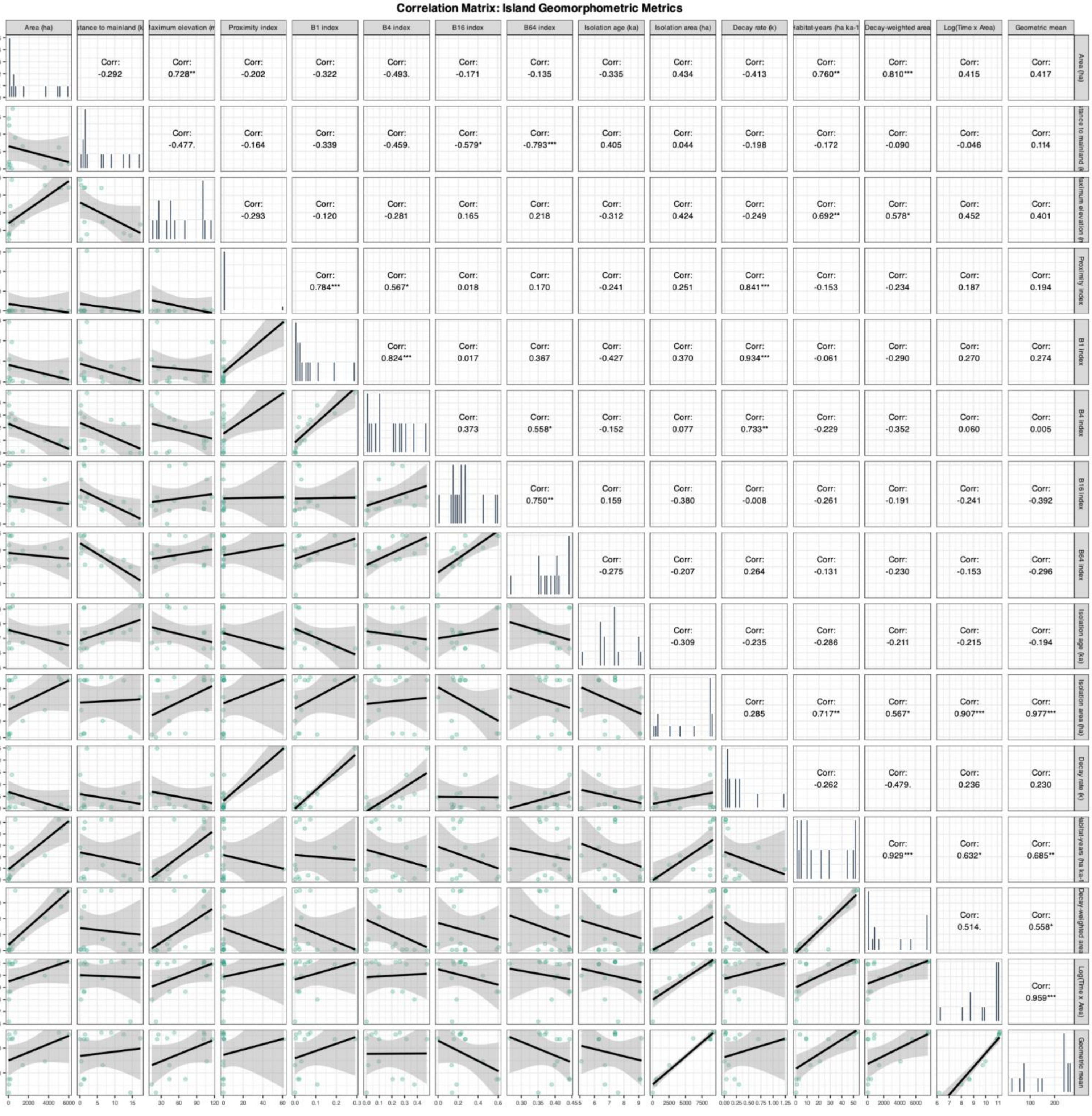

**Fig. S3. Correlation matrix of all island geomorphometrics, modern, historical and combined.** Significant relationships reveal autocorrelations that may obscure detection of explanatory mechanisms.

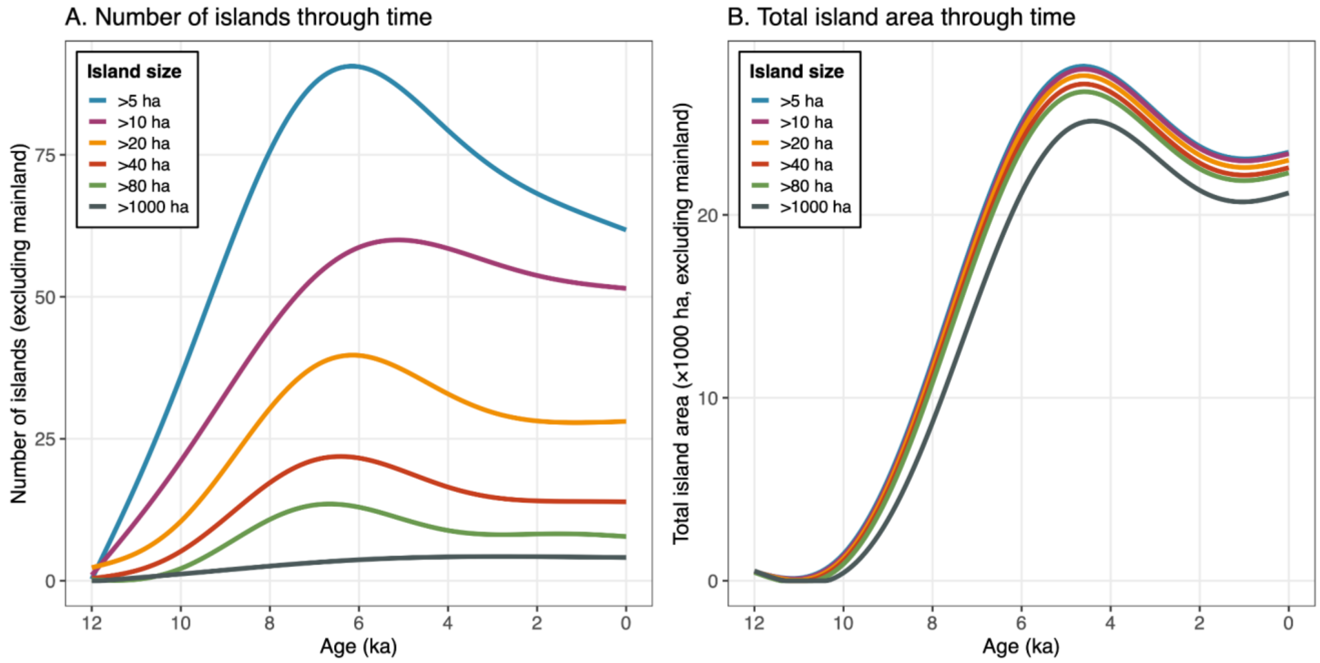

**Fig. S4. The effect of island threshold size** on estimates of number of islands (A) and total island area (B).

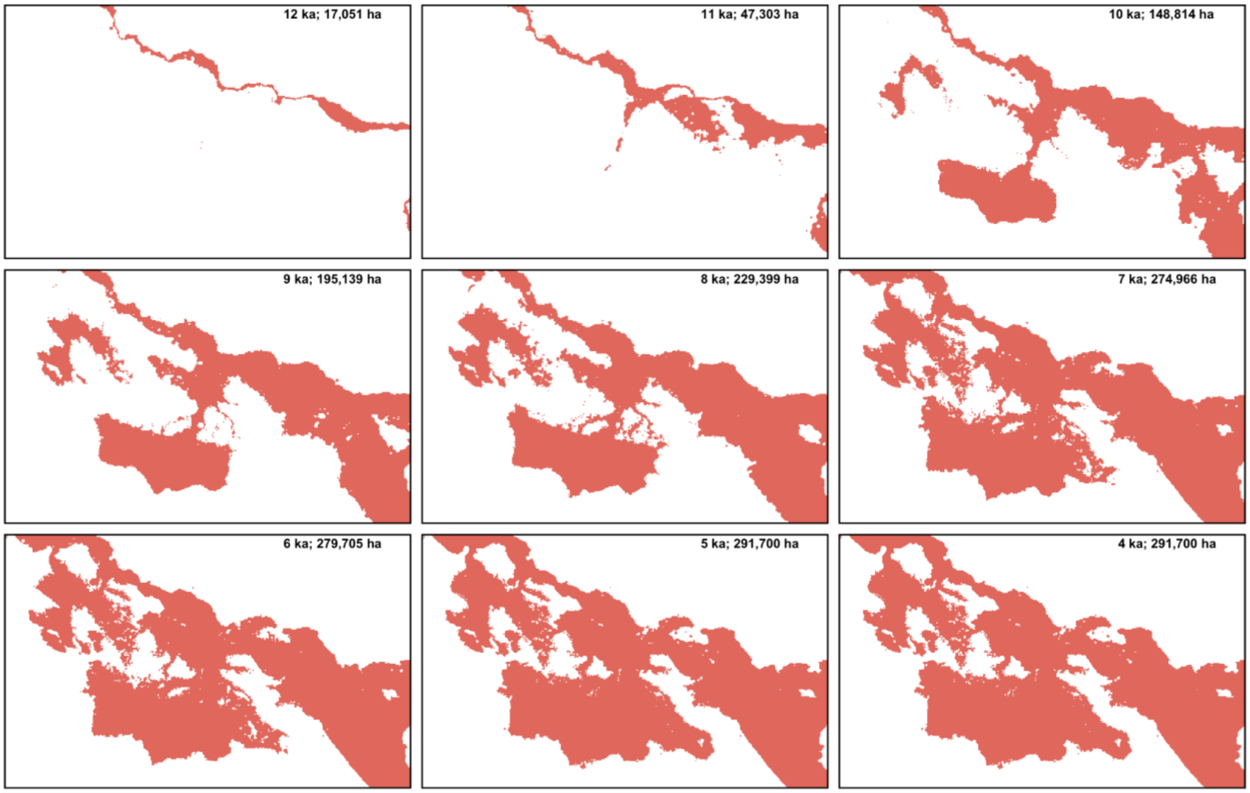

**Fig. S5. Estimated extents of shallow marine habitat (defined here at 0-50m depth) through time.** Red areas represent seafloor at 0-50m depth below sea level, corresponding to the depth range that harbour a large proportion of marine biodiversity.

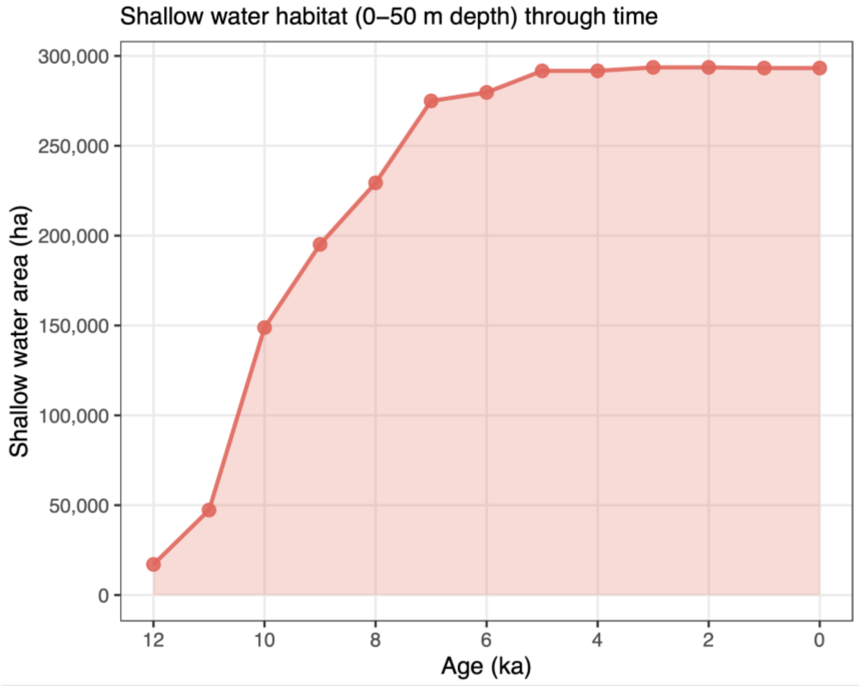

**Figure S6. Area of shallow water seafloor (0–50 m depth) in Bocas del Toro over the last 12 kyr.** This area represents the available habitat for the majority of tropical marine diversity. The 0–50 m depth range encompasses the photic zone where most reef-building corals, seagrasses, and macroalgal communities develop.

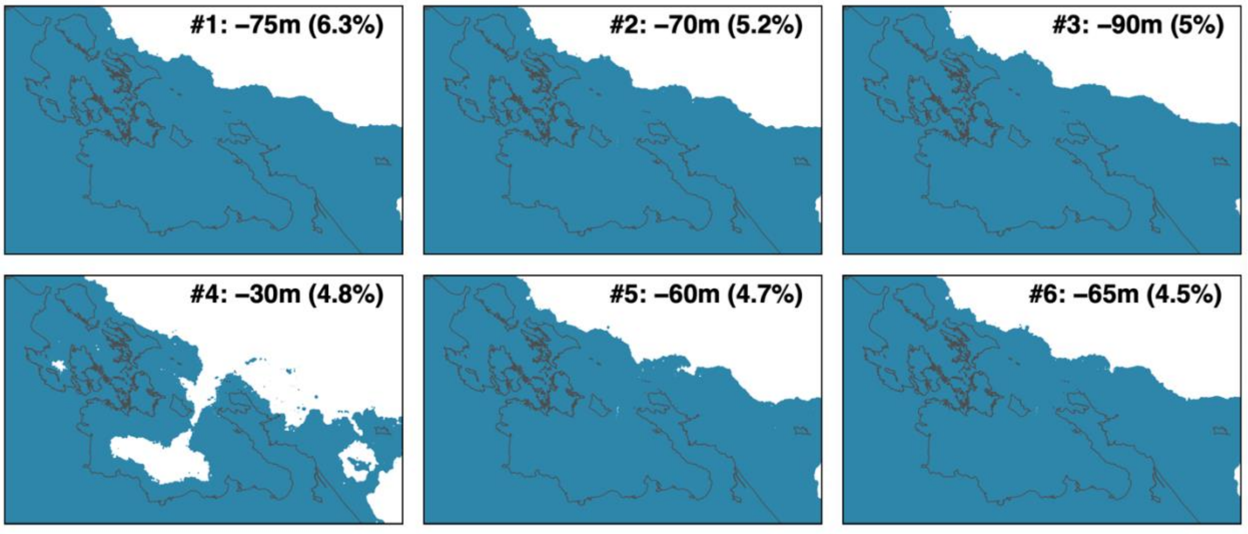

**Figure S7. Land configurations for the six most frequently occurring sea level states over the last 1 Myr.** Maps show reconstructed land area for each of the six most common sea level states, ranked by cumulative duration and binned into 5 m intervals (±2.5 m). Percentage values indicate the proportion of the last million years spent at each state. Modern coastline outlined by thin dark grey line. Together these six states account for over 31% of the past million years; none produces an archipelago resembling the modern configuration.

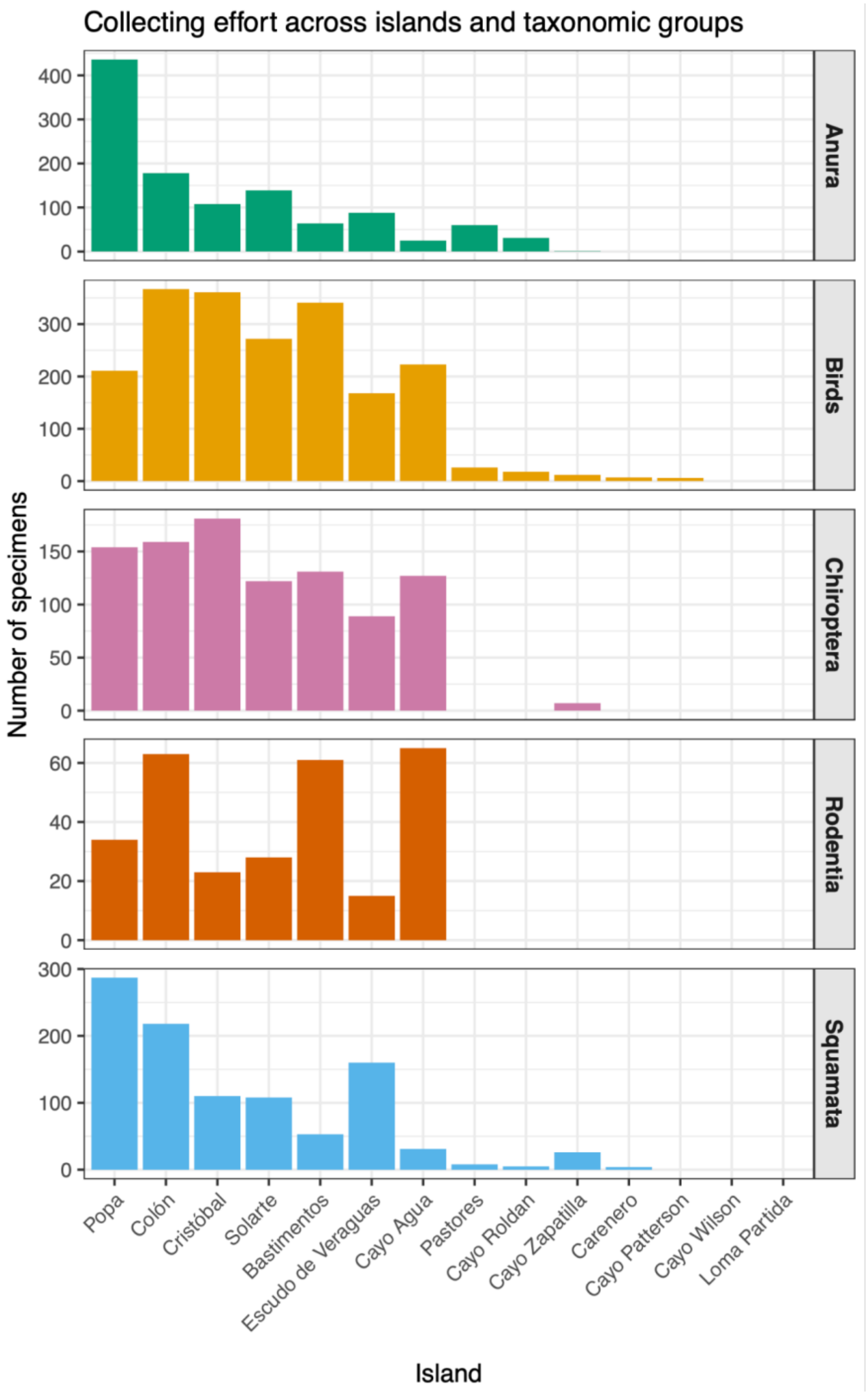

**Figure S8. Variation in museum collecting effort across islands and taxonomic groups.** Bar charts show specimen counts per island for five terrestrial vertebrate groups. Islands are ordered by total specimens (highest to lowest, left to right). Collecting effort varies 10–100× among islands. Totals for all islands: 5,411 specimens, 273 species, 14 islands.

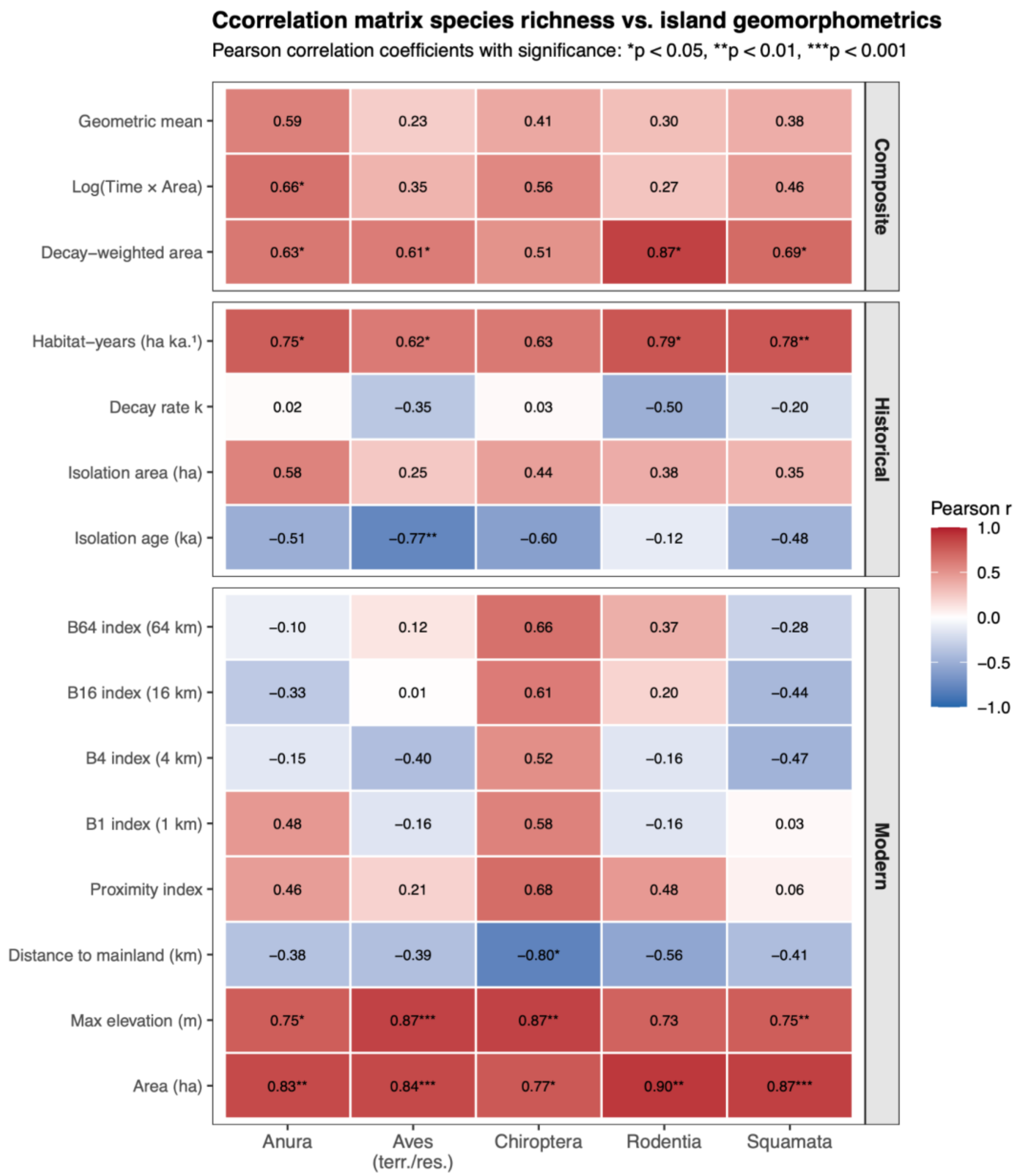

**Figure S9. Complete correlation matrix of species richness versus all 15 island geomorphometric metrics for five terrestrial vertebrate groups.** Heatmap shows Pearson correlation coefficients (r) between species richness and modern metrics (island area, distance to mainland, maximum elevation, proximity index, buffer zone indices B1–B64), historical metrics (isolation age, isolation area, decay rate, cumulative habitat-years), and composite metrics (decay-weighted area, log[time × area], geometric mean). Significance levels: *$p < 0.05$, **$p < 0.01$, ***$p < 0.001$. This expanded heatmap complements the summary shown in Fig. 7A, which presents only the six key metrics. Note the consistently strong positive correlations with area and elevation across all groups, and the generally weak or negative correlations with buffer zone isolation indices, reflecting the structural confound between island size and surrounding landmass in continental shelf archipelagos (see main text).
